# Blue Light Promotes Root Iron Acquisition via a Shoot CRY–HY5 Signaling Module in Arabidopsis

**DOI:** 10.1101/2025.11.02.686065

**Authors:** Pooja Jakhar, Samriti Mankotia, Shubham Ladhwal, Nisha Bodh, Krishna Dhanawat, Ajay K. Pandey, Santosh B. Satbhai

## Abstract

- Iron (Fe) is an essential element for almost all living organisms and its deficiency severely impacts crop productivity and human health. While the mechanisms underlying iron uptake and accumulation are well studied, the role of light in modulating iron homeostasis remains poorly understood. In this study, we report that blue light enhances iron accumulation thereby positively regulating iron homeostasis in *Arabidopsis thaliana*.
- Under iron-deficient conditions, root elongation is specifically promoted by blue light, not red light. We further revealed that, light perceived by shoots is required for optimum iron accumulation and for iron deficiency induced increase in root length.
- The transcription factor HY5 (Elongated Hypocotyl 5), mediates these blue light-driven responses. The *hy5* mutant showed reduced root elongation under -Fe conditions as well as iron accumulation under blue light as compared to wild-type plants.
- Further we report that CRY1 and CRY2 are the photoreceptor that acts as a connecting link between blue light and HY5 to regulate iron uptake in Arabidopsis.
- Taken together, our work reveals new regulatory mechanism and offers potential insights into optimizing light conditions to enhance iron accumulation and improve the nutritional quality of crops.

## Introduction

Plants, being sessile organisms, are continually exposed to various environmental stresses and have evolved intricate physiological, molecular and biochemical strategies to adapt and thrive under these changing conditions. Iron (Fe) deficiency is one of the most widespread nutritional stress affecting both plants and humans. While Fe toxicity can be detrimental, Fe deficiency due to its limited bioavailability in aerobic soils remains a global concern. Iron plays a crucial role in various fundamental processes, including chlorophyll synthesis, photosynthesis, respiration and redox homeostasis (Kobayashi & Nishizawa, 2012). Despite being one of the most abundant elements in Earth’s crust, Fe is mostly present in insoluble ferric (Fe^3+^) form under aerobic and alkaline conditions, making it inaccessible to plants (Darbani et al., 2013). Importantly, Fe deficiency not only affects plant productivity, it also contributes to Fe malnutrition in humans. Therefore, enhancing Fe accumulation and acquisition in plants is of agronomic and nutritional importance, which can be a step towards combating anemia, and improve micronutrient content in crops.

To overcome Fe limitation, plants have evolved two different strategies. Grasses employ a chelation-based strategy (Strategy II), in which siderophores belonging to mugineic acid (MA) family are released in the rhizosphere, which chelates Fe^+3^ and the Fe (3)-MA complexes are taken up by roots (Kobayashi et al., 2019; Römheld & Marschner, 1986; Takahashi et al., 2001). In contrast, dicots and non-graminaceous monocots such as *Arabidopsis thaliana* (Arabidopsis) utilize a reduction-based mechanism (Strategy I), involving acidification of the rhizosphere by the plasma membrane H⁺-ATPase AHA2, secretion of Fe-mobilizing coumarins via the ABCG transporter PDR9, and subsequent reduction of Fe³⁺ to Fe²⁺ by FERRIC REDUCTION OXIDASE 2 (FRO2) before transport into root cells through the Fe²⁺ transporter IRT1 (Connolly et al., 2003; Eide et al., 1996; Fourcroy et al., 2014; Henriques et al., 2002; Robinson et al., 1999; Santi & Schmidt, 2009; Vert et al., 2002). *AHA2*, *FRO2* and *IRT1* are the major iron uptake genes, regulated via an intricate machinery involving various transcription factors (Ivanov et al., 2012). In response to iron deficiency, expression of key Fe acquisition genes *IRT1, FRO2* is tightly regulated by the FIT (FER like Iron deficiency induced transcription factor) and its interaction with bHLH Ib subgroup transcription factors (basic helix-loop-helix), bHLH38, bHLH39, bHLH100 and bHLH101 (Colangelo & Guerinot, 2004; Maurer et al., 2014; Wang et al., 2013; Yuan et al., 2008). Upstream of this module, bHLH121/URI (Upstream Regulator of IRT1) and subgroup IVc bHLH transcription factors (bHLH34, bHLH104, bHLH105/ILR3, and bHLH115) form a hierarchical regulatory network that modulates Fe deficiency responses (Kim et al., 2019; Li, Lei, et al., 2021; Liang et al., 2017; J. Zhang et al., 2015). Moreover, the E3 ubiquitin ligase BRUTUS (BTS) and its homologues BTSL1 and BTSL2 act as negative regulators of Fe homeostasis by targeting bHLH IVc transcription factors and URI for degradation under Fe-sufficient conditions (Hindt et al., 2017; Long et al., 2010a; Selote et al., 2015).

Recent reports have revealed that additional layer of regulation via Fe Uptake–Inducing Peptides (FEPs/IMAs), which competitively inhibit BTS–bHLH interactions, thereby enhancing Fe deficiency responses (Grillet et al., 2018; Hirayama et al., 2018; Li, Lei, et al., 2021; Lichtblau et al., 2022). Furthermore, bHLH105(ILR3) forms repressive heterodimers with the POPEYE (PYE) or bHLH47, which restricts expression of genes involved in Fe assimilation, transport, and storage to prevent Fe overload (Long et al., 2010; Tissot et al., 2019). Interestingly, PYE has recently been shown to interact with ELONGATED HYPOCOTYL 5 (HY5), a central transcriptional regulator of light signaling, linking Fe homeostasis to light perception (Mankotia et al., 2025). Recent evidence reveals an additional light-dependent regulatory mechanism whereby FIT forms blue light–induced subnuclear condensates (“FIT nuclear bodies”) that enhance its activity through interactions with bHLH Ib partners (Trofimov et al., 2024), suggesting that light cues may directly modulate Fe uptake mechanisms.

Light is a crucial environmental cue for plant growth and development, affecting various components of plant life, including seed germination, de-etiolation, organogenesis and flowering (Quail, 2002). Light has increasingly been recognized as an important regulator of root-associated nutrient dynamics. Shoot light perception has been shown to affect root uptake of several essential nutrients, including nitrogen, phosphorus, sulfur, and copper (Chen et al., 2021; X. Chen et al., 2016; Gao et al., 2021; Sakuraba et al., 2018; X. Wang et al., 2023; H. Zhang et al., 2014). Such light-mediated signaling is coordinated by photoreceptors (phytochromes, cryptochromes, phototropins, and UVR8) that activate diverse transcriptional networks via bZIP, bHLH, and MYB transcription factors (Quail, 2002).

Recent studies have revealed a direct role for HY5 in Fe homeostasis. In tomato, red light–induced accumulation of *SlHY5* upregulates *SlFER*, a positive regulator of Fe uptake (Guo et al., 2021a). In Arabidopsis, HY5 directly binds to promoters of Fe uptake gene *FRO2*, while repressing negative regulators *BTS* and *PYE*, thereby promoting Fe acquisition under Fe deficiency (Mankotia et al., 2023). Blue light has also been shown to enhance Fe reductase activity and increase *IRT1* and *FRO2* expression (Trofimov et al., 2024), suggesting that blue light–mediated HY5 activation may coordinate Fe uptake. Together, these findings suggest that photoreceptor-mediated stabilization of HY5, may represent a central mechanism integrating light signaling and Fe homeostasis.

In this study, we investigated the role of blue light and cryptochrome signaling in regulating Fe uptake and homeostasis in *Arabidopsis thaliana*. We showed that blue light enhances Fe deficiency responses through CRY1/CRY2-mediated activation of HY5. Our findings reveal that the CRY–HY5 signaling module plays an important role in promoting Fe uptake and accumulation, connecting aboveground light perception with belowground nutrient acquisition. Our findings offer new insights into the integration of light signaling with nutrient acquisition pathways and highlight the potential of optimizing light conditions, particularly in controlled growth environment, as a promising strategy to enhance Fe uptake and improve nutritional quality of crops.

## Material and methods

### Plant materials and growth conditions

In this study, *Arabidopsis thaliana* ecotype Columbia (Col-0) was used as wild type (WT) for all the experiments, unless mentioned otherwise. The mutants and transgenic lines used for this work are *hy5* (SALK_056405), phyB-9 (Reed et al., 1993), *phot1-5 phot2-1*(Vandenbussche et al., 2014), *pHY5:HY5: YFP/hy5* (Wang et al., 2021), *cry1-304, cry2-1, cry1cry2(cry1-304cry2-1*) (Mockler et al., 1999), *cop1-4* (Datta et al., 2005), *hy5-215cop1-4* (Yadukrishnan et al., 2020), p*IRT1*:*GUS*/Col-0 (Blum et al., 2014) and p*IRT1*:*GUS*/*hy5* (Mankotia et al., 2023). Additionally we used, *hy5, hyh* and *hy5/hyh* in the WS background and Wassilewskija (WS-0) ecotype as previously described (Binkert et al., 2014).

*hy5* single mutant was crossed with *cry1-304, cry2-1* to obtain double mutants and *cry1cry2* to obtain triple mutants. Initial screening was done based on phenotype, followed by confirmation with genotyping for *hy5* mutant, using primers mentioned in Table 1, and by immunoblotting using anti-CRY1 (Agrisera, AS16-3933) and anti-CRY2 (Agrisera, AS18-4221) antibodies. The seeds were surface sterilized using 5% sodium hypochlorite for 2 minutes, followed by a 2 minutes wash with 70% ethanol and 3-4 washes of autoclaved water. Seeds were stratified at 4°C for 3/4 days. The plants were grown on square plates containing 50ml of ½ MS (Murashige and Skoog) media (Caisson Labs) containing Fe(+Fe) or iron free ½ MS media (-Fe) (Caisson Labs) for iron limiting conditions. Ferrozine (3-(2-pyridyl)-5,6-diphenyl-1,2,4-triazine sulfonate) (Sigma-Aldrich, Bangalore, India) was added to iron free media at 300 µM concentration to create iron deficiency, wherever required. The media was supplemented with 1% sucrose and 1% agar and pH was kept 5.74, adjusted using 1M KOH. The plates were kept vertically in growth chamber under long day conditions, 16 hours light and 8 hours dark at 22°C with 50 % humidity and different light conditions in ARA-LAB plant chamber. The light intensity for white light was kept 110/120 μmol m^-2^ s^-1^, for blue light 70 μmol m^-2^ s^-1^ and red light 55 μmol m^-2^ s^-1^. For soil, a mixture of soilrite, perlite and compost (3:1:0.5) was used to grow plants.

### Phenotypic analysis

Seeds were directly plated on ½ MS (Murashige and Skoog) media (Caisson Labs) containing Fe (+Fe) and on iron free ½ MS media (-Fe) (Caisson Labs) for iron limiting conditions. The plates were kept vertically in growth chamber under different light conditions, white, blue and red respectively. For transfer experiments, plates were kept in white light for 5 days and then transferred to different light conditions for another 5 days. After 10 days, the plates were scanned using Epson Perfection V600 scanner at 1200 dpi resolution and root length was measured using ImageJ 1.52a software (National Institute of Health).

### Chlorophyll content measurement

The total chlorophyll content was measured using seedlings grown on +Fe and -Fe for 10 days under different light conditions. 5 shoots were pooled as one sample. The samples were weighed and incubated in a 1ml solution of 80% acetone for 24 hours in dark, followed by measurement of absorbance at A645 nm and A663 nm. The total chlorophyll content was quantified using the formula, Chlorophyll content (mg/gFW) = (20.3 × A645 ± 8.04 × A663) × V/W × 10^3^, as described previously (Aono et al., 1993).

### Ferric chelate reductase (FCR) assay

The seedlings were grown on +Fe for 5 days and then transferred to +Fe and -Fe+300µM FZ media for 3 days under different lights. The roots from ∼10 seedlings were pooled as one sample, weighed and incubated for one hour in 700 µl of assay solution composed of 0.1 mM Fe (III)-EDTA and 0.3 mM ferrozine. The activity of FRO2 leads to formation of a purplish compound, which is measured spectrophotometrically by taking absorbance at 562 nm. The activity was calculated using formula [nmol Fe(Ⅱ) per g FW per hr = (A/28.6) × V/Root FW] (Yi & Guerinot, 1996).

### Perls’ and Perls’ DAB staining for iron content

Seedlings grown on +Fe for 5 days under different light conditions were vacuum infiltrated for 30 minutes in a solution consisting of 2% (v/v) HCl and 2% (w/v) K-ferrocyanide (Perls’ solution). The seedlings were then washed three times with water followed by a single wash of 70% ethanol. The seedlings were observed under a NIKON ECLIPSE Ni U microscope and images were captured. The intensity of blue color depicts comparative Fe^3+^ content.

For total iron staining (Fe^3+^ and Fe^2+^), DAB(Diaminobenzidine) intensification was done. The Perl’s stained seedlings were incubated in a methanol solution, supplemented with 10 mm Na-azide and 0.3% (v/v) H2O2, for one hour at room temperature. The seedlings were then washed with 100 mm Na phosphate buffer (pH 7.4) followed by incubation for 1.5 minutes in DAB solution composed of 0.025% (w/v) DAB and 0.005% (v/v) H2O2) in 100 mm Na phosphate buffer (pH 7.4). The seedlings were washed with 70% ethanol and observed under a NIKON ECLIPSE Ni U microscope and images were captured.

### Grafting

Reciprocal grafting was carried out as per a previously described protocol(Marsch-Martínez et al., 2013). Briefly, WT, *hy5* and *cry1cry2* seedlings were grown on ½ MS media Iron sufficient media (+Fe) (Caisson Labs) supplemented with 1% sucrose and 1% agar for 5days under white light. The cotyledons were removed and the scion and stalk were separated and moved to fresh ½ MS Iron sufficient media (+Fe) (Caisson Labs) plates. The scion and stalk were aligned as per their respective combinations. After 5days of grafting, the successfully grafted pairs were used for Perls’ staining.

### Iron content quantification

A spectrophotometric method was used for iron content quantification (Gautam et al., 2022). The shoots and roots from samples grown under different lights were separated and ∼30 shoots/roots were pooled as one biological replicate and three such replicates were used. The samples were dried for 2 days in an oven at 60°C followed by their dry weight measurement. The samples were then digested with 65% (v/v) HNO3 at 95°C for 6 h, followed with addition of 30% (v/v) H2O2 at 56°C for 2 h.

An assay solution composed of 1 mM bathophenanthroline disulfonate (BPDS), 0.6 M sodium acetate and 0.48 M hydroxylamine hydrochloride was prepared. A standard curve was made using standards containing different known concentrations of FeCl3. Assay solution when mixed with sample containing iron, forms a pink coloured Fe^2+^–BPDS_3_ complex, which can be quantified by taking absorbance at 535nm. The iron content was quantified using the equation obtained from the standard curve, followed by normalization with dry weight for each sample.

### GUS staining

The *pIRT1:GUS*/Col-0 and *pIRT1:GUS/hy5* seedlings were grown on +Fe and -Fe for 5 days or transferred to -Fe+300µM FZ for 3 days under different light conditions, followed by vacuum infiltration with staining solution having 100 mM Na_2_HPO_4_, 2 mM K_3_[Fe(CN)_6_], 2 mM K_4_[Fe(CN)_6_], Triton X-100, 100 mM NaH_2_PO_4_, and 2 mM 5-bromo-4-chloro-3-indolyl glucuronide for 25 minutes. Subsequently the seedlings were incubated at 37°C for 2.5 hours, washed with 70% ethanol and imaged using NIKON ECLIPSE Ni U microscope.

### Confocal microscopy

The p*HY5*:*HY5*:GFP/*hy5* and p*FRO2*:*FRO2*:mCherry/*frd1*seedlings were used for confocal imaging. The seedlings grown on +Fe and -Fe for 5 days or transferred to -Fe+300µM FZ for 3 days under different light conditions. Seedlings were stained with PI (Propidium Iodide) for 2 minutes and rinsed with water. For GFP, a 488-nm laser was used for excitation, and propidium iodide was excited with 561-nm laser, and emission spectra were collected at 500–530 nm and 600–650 nm, respectively. For mCherry the excitation and emission were at 587nm and 610 respectively. Imaging was done in Z-stack mode with 2 µm step size using a SP8 upright confocal microscope (Leica). The Z-stack was processed to maximal Z-projection (Merge, PI and GFP/mCherry). Scale bar: 100µm. Fiji® software Macro was used to quantify raw integrated density of the signal.

### RNA Extraction and qRT-PCR

The seedlings were grown on +Fe for 5 days and then transferred to +Fe and -Fe+300µM FZ media for 3 days under different lights. Total RNA was extracted using root samples using Plant RNeasy kit (Qiagen) following manufacturer’s protocol, followed by digestion of ∼2 µg of extracted RNA with DNaseI. Around ∼1 µg of DNase treated RNA was used for cDNA synthesis using RevertAid^TM^ First Strand cDNA Synthesis Kit (Thermofisher). The concentrated cDNA so formed was diluted using nuclease free water to get a working solution of 50ng/µl concentration, which was used as a template for qRT-PCR, performed with TB Green^TM^ Premix Ex (Takara) using a Qiagen QIAquant RT PCR system. TUBULIN was used as an internal control and relative expression levels were quantified using ΔΔCt method. The primers used for qRT-PCR are listed in Table 1.

### Immunoblotting

Proteins were isolated from the seedlings were grown on +Fe for 5 days and then transferred to +Fe and -Fe+300µM FZ media for 3 days under different lights. The tissue, root/shoot/whole seedling, depending on experiment, was harvested and crushed using liquid nitrogen and 300µl of protein extractions buffer composed of Tris–HCl (125 mM) (pH 7.5), 5% β-mercaptoethanol, 0.5% SDS, 10% glycerol, 1% Tween-20, 2 mM EDTA, 1% protease arrest inhibitors (G Bioscience) and 1 mM PMSF followed with TCA precipitation. The concentration of each sample was determined using Bradford assay. Around ∼25 µg protein from each sample was heated at 75°C for 15 minutes, followed by separation using 12% SDS-PAGE. The proteins were then transferred to PVDF (polyvinylidene difluoride) membrane by wet transfer method. The protein of interest was detected using corresponding antibody. HY5-GFP was detected with 1:7000 dilution of anti-GFP primary antibody (Abcam, AB290), IRT1 with 1:7000 anti-IRT1 primary antibody (Agrisera, AS111780), HY5 using a 1:1000 dilution of primary antibody, anti-HY5 (Agrisera, AS12-1867), CRY1 and CRY2 using 1:1000 dilution of anti-CRY1 (Agrisera, AS16-3933) and anti-CRY2 (Agrisera, AS18-4221) respectively. All the primary antibodies were diluted in 5% BSA in TBST. After washing, the membrane was incubated with a HRP-conjugated secondary antibody (Sigma-Aldrich; AP307P) diluted 1:15 000 in 5% non-fat milk in TBST. The membrane was again washed using TBST followed by detection using ECL select HRP substrate (Bio-Rad; #170-5060). The same membrane was stripped using protein stripping buffer, followed by washes with 1x TBST. The membrane was again incubated with 1:7000 dilution of anti-Tubulin antibody as the internal control (CST 3837S Alpha-Tubulin mouse mAb), in 5% non-fat milk in TBST followed by secondary antibody treatment with 1:20000 dilution of Anti-Mouse (Sigma A4416) in 5% non-fat milk in TBST. The membrane was redeveloped using ECL select HRP substrate (Bio-Rad; #170-5060).

The intensity of the HY5/IRT1/CRY1/CRY2 band and Tubulin band was quantified using IMAGEJ software (US National Institutes of Health, Bethesda, MD, USA), and the HY5/IRT1/CRY1/CRY2 values were divided by the Tubulin values for each sample. The HY5/IRT1/CRY1/CRY2 level in the white light Wild type sample, grown on iron sufficient media, was set to 1 from these ratios and the relative values for different conditions were calculated. These relative values are shown above their respective bands.

### Statistical analysis

The graphs were plotted using GraphPad PRISM10. Significance was calculated using Student’s t-test or one-way ANOVA followed by a post hoc Tukey HSD test. Different letters were used to (a, b, c) indicate significant differences. Error bars represented the ± SEM. A p-value ≤ 0.05 was considered statistically significant.

## Results

### Light sensing in shoots controls iron accumulation and root elongation

Previously it has been reported that iron absorption rate decreases in absence of light (Guo et al., 2021a). To assess the role of light in Fe accumulation, plates containing wild type (WT) seedlings were covered with aluminium foil to induce darkness. Fe accumulation was evaluated using Perl’s and Perls’ DAB staining and we observed severely impaired iron accumulation in seedlings grown in the dark (Fig. 1a). Subsequently, WT seedlings were grown under standard long day conditions (16 hr light/8 hr dark) for five days, followed by transfer to continuous darkness for three days. This transfer assay also resulted in a strong reduction of iron content in the roots (Fig. 1a). These results confirmed that plants fail to uptake iron under dark conditions and require light for optimum iron acquisition and accumulation thus supporting that iron uptake is light-dependent. Under natural conditions, light is predominantly perceived by the shoots, however, when seedlings are grown vertically on agar plates in a growth chamber, both shoots and roots are exposed to light. To differentiate tissue specific role of light, seedlings were grown on iron-sufficient media under four different light exposure conditions: (1) full light exposure (shoot and root), (2) shoot specific darkness (upper ∼2 cm of plates covered with foil), (3) root specific darkness (only ∼2 cm upper region exposed), and (4) complete darkness (entire plate covered with foil). Next, Fe accumulation was observed using Perl’s and Perls’ DAB staining. The results indicated that Fe accumulation is not affected when only the roots were kept in darkness, whereas significant reduction was observed when shoots were kept in the dark as compared to uncovered seedlings. These results indicated that light sensing in the shoot is vital for iron accumulation and likely a light-dependent signaling controls Fe homeostasis (Fig. 1b).

**Fig. 1.**
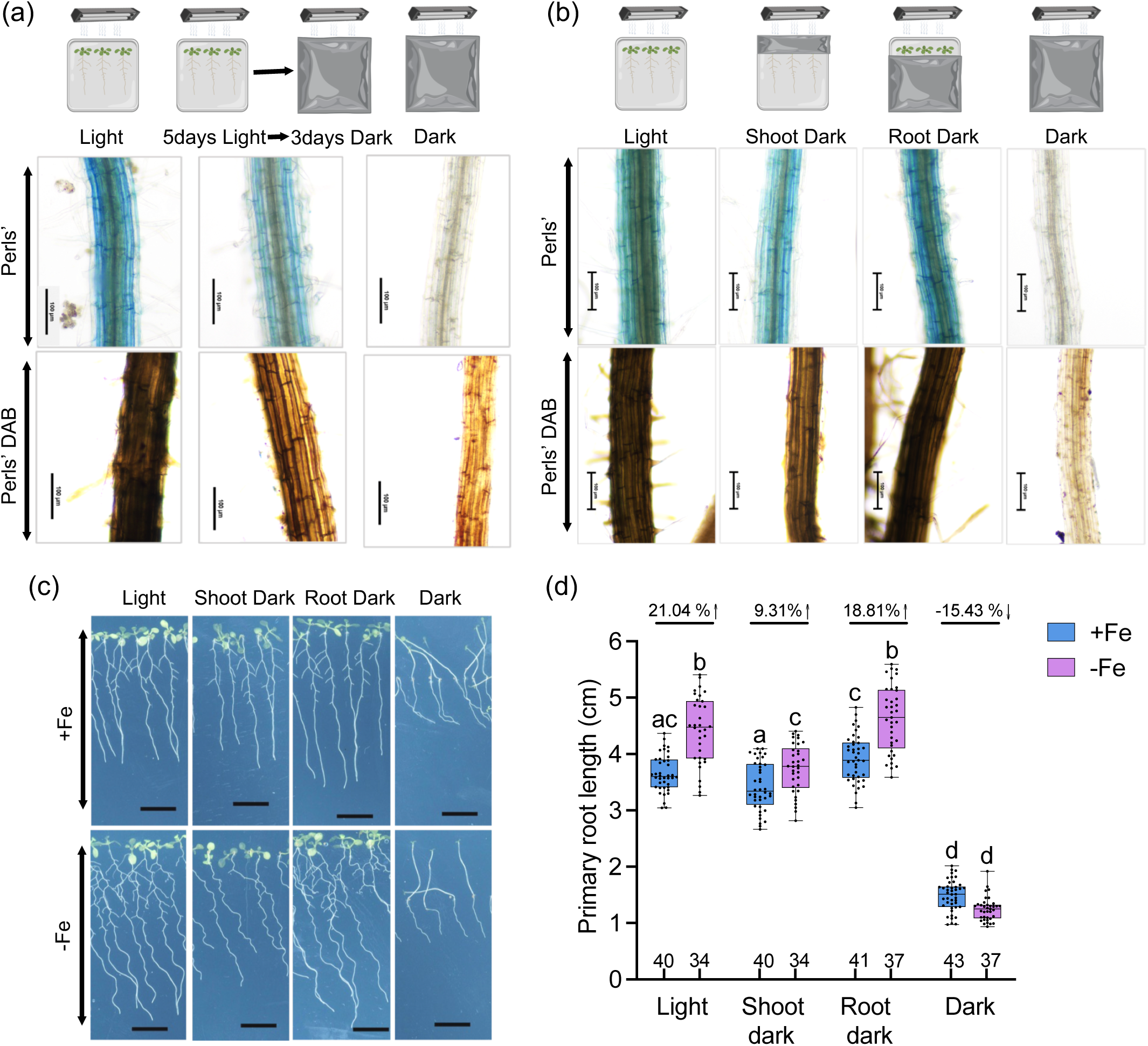
Light sensing in shoots controls iron accumulation and root elongation. (a) Perls’ and Perls’ DAB stained maturation zone of WT plants grown on Fe-sufficient (+Fe) media under normal long day, complete dark (covered with foil) and covered with foil for 3 days after 5 days of normal long day treatment. Scale bar: 100 µM. (b) Perls’ and Perls’ DAB stained maturation zone of WT plants grown on Fe-sufficient (+Fe) media under normal long day with full light exposure (shoot and root), shoot specific darkness (upper ∼2 cm of plates covered with foil), root specific darkness (only ∼2 cm upper region exposed), and complete darkness for 5 days. Scale bar: 100 µM. (c) Phenotype of WT plants grown on Fe-sufficient(+Fe) and Fe-deficient (-Fe) media under normal long day, complete dark (covered with foil), shoot dark (shoot covered with foil) and root dark (roots covered with foil) for 10 days. Scale bar:1 cm. (d) Boxplot of root length of WT plants grown on +Fe and -Fe media under normal long day, complete dark (covered with foil), shoot dark (shoot covered with foil) and root dark (roots covered with foil) for 10 days. Means within each condition with the same letter are not significantly different according to one-way ANOVA followed by post hoc Tukey test, P < 0.05. using GraphPad PRISM 10. In (a-b) :-Images showing experimental set up made using Biorender.

Previous studies have reported that root length increases under iron deficiency conditions (Gruber et al., 2013; Mankotia et al., 2023). Consistent with these findings, we observed ∼18.81% increase in root length under Fe deficient conditions when only the roots were covered with foil, which is comparable to ∼21.04% increase when seedlings were grown under normal light conditions. In contrast, when shoots were covered, only a 9.31% increase was seen which might be due to a slight percolation of light from the sides and occasional drooping of shoots below the foil-covered region. Notably, under complete darkness, roots were ∼15.43% shorter under iron deficient conditions compared to iron sufficient conditions (Fig. 1c-d). This result further strengthens the conclusion that light is a critical factor not only for the iron accumulation but also for promoting root growth in response to iron deficiency.

### Blue light is required for optimum iron uptake and iron deficiency induced increase in root length

The different lights in sunlight spectra like blue, red, far-red and UV are known to be involved in regulation of plant growth and development and recent studies have highlighted their roles in nutrient homeostasis (Gao et al., 2021; Guo et al., 2021a; Wang et al., 2023). To investigate the specific roles of blue and red light on iron homeostasis, Arabidopsis seedlings were grown under white light (110 μmol m^-2^ s^-1^), blue light (70 μmol m^-2^ s^-1^) and red light (55 μmol m^-2^ s^-1^) in long day conditions (16 h light / 8 h dark). The seedlings grown under white light showed a ∼25.54% increase in root length under iron deficient conditions as compared to iron sufficient conditions. Notably, the blue light grown seedlings showed an even greater response with root length increasing by ∼49.73% nearly double of that observed under white light (Fig 2 a-b). In contrast, seedlings exposed to red light did not show any significant change in root length (∼0.46% only) under Fe deficient conditions (Fig. 2a-b). Perls’ and Perls’ DAB staining further revealed enhanced Fe accumulation in blue light grown seedlings, as indicated by more intense staining compared to white light (Fig. 2c). Conversely, red light grown seedlings showed minimal or no staining, suggesting that iron uptake is hindered in red light (Fig. 2c). Moreover, quantitative analysis showed that the iron content in blue light grown seedlings was ∼15% higher in shoots and ∼20% higher in roots as compared to those grown under white light (Fig 2d). In addition, the iron content was substantially reduced in seedlings grown under red light (∼56% in shoots and ∼89% in roots) as compared to the seedlings grown under white light (Fig 2 d).

**Fig. 2.**
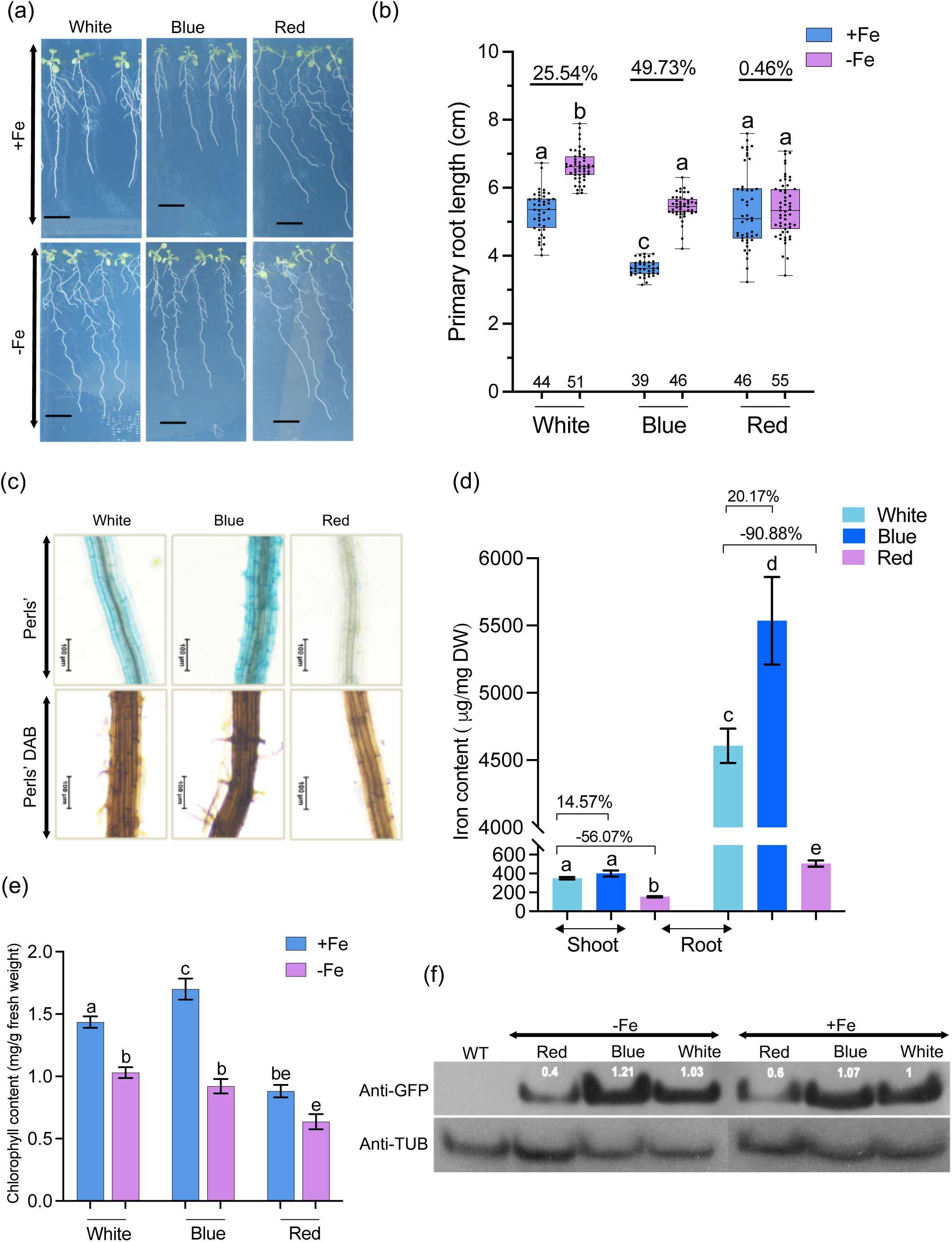
Blue light is required for iron accumulation and root growth under iron deficiency. (a) Phenotype of WT plants grown on +Fe and -Fe media under white, blue and red light for 10 days. Scale bar: 1 cm. (b) Boxplot of root length of WT plants grown on +Fe and -Fe media under white, blue and red light for 10 days. * indicate significant difference according to one-way ANOVA followed by post hoc Tukey test, P < 0.05 using GraphPad PRISM 10. (c) Perls’ and Perls’ DAB stained maturation zone of WT plants grown on +Fe media under white, blue and red light for 5 days of normal long day cycle. Scale bar: 100 µM. (d) Iron content quantified spectrophotometrically in shoots and roots of WT plants grown on +Fe media under white, blue and red light for 15 days. The data represent average of three biological replicates, with three technical replicates of each. Each biological replicate consisted of ∼30 roots/shoots. Error bars represent ±SEM. DW - dry weight. Different letters (a, b, c, d) indicate significant differences, determined by one-way ANOVA followed by a post hoc Tukey Test (P≤0.05) using GraphPad PRISM 10. (e) Total chlorophyll content in shoots of WT plants grown on +Fe and - Fe media under white, blue and red light for 10 days. Each biological replicate consisted of 5 shoots. Error bars represent ±SEM. Different letters (a, b, c, d) indicate significant differences, determined by one-way ANOVA followed by a post hoc Tukey Test (P≤0.05) using GraphPad PRISM 10. (f) Western Blot showing HY5 protein levels in *pHY5:HY5:GFP/hy5* seedlings grown on +Fe media for five days and then transferred to +Fe and –Fe (+300 µM Fz) media for 3 days, under different light conditions. WT-Negative Control, Tubulin - Internal Control. The values above each band depicts the intensities after normalization with internal control, followed by normalization using White light +Fe as a control.

Furthermore, the chlorophyll content was higher in seedlings grown under blue light, consistent with higher iron content, while a severe reduction was seen in red light grown seedlings (Fig. 2e). Time-course experiments in which white light grown seedlings were transferred to blue or red light further supported these findings, where iron accumulation significantly increased after 72 hours under blue light, whereas red light caused a reduction in iron levels as early as 24 hours post-transfer (Fig. S1). (Fig. S1). Collectively, these results demonstrate that blue light promote iron accumulation and positively regulates the iron deficiency induced increase in root length while red light did not enhance accumulation and had no effect on root growth as seen in white or blue light conditions.

Previous studies have established that HY5 functions as key regulator of iron homeostasis via directly regulating the expression of important iron homeostasis genes, including *FRO2, BTS, PYE* and *NAS4* (Mankotia et al., 2023, 2025). HY5 is also a well-known central player in photomorphogenesis and nutrient signaling (Chen et al., 2016; Gangappa & Botto, 2016; Gao et al., 2021; Mankotia et al., 2023). To investigate whether HY5 protein accumulation is influenced by light quality under iron-sufficient and iron-deficient conditions, we assessed HY5 levels using immunoblot analysis. The blue light grown seedlings accumulated slightly higher amount of HY5 protein especially under -Fe conditions, as opposed to red light conditions where amount of HY5 protein was reduced (Fig. 2f). These findings suggest that blue light not only promotes iron acquisition but also enhances HY5 protein stability or accumulation under iron-limiting conditions.

### Blue light specific regulation of iron homeostasis is HY5 mediated

HY5, a key regulator of photomorphogenesis, thermotolerance, and nutrient signaling, links light perception with nutrient homeostasis.(X. Chen et al., 2016; Gangappa & Botto, 2016; Gao et al., 2021a; Guo et al., 2021b; Li, Shi, et al., 2021; P. Liu et al., 2025; S. Liu et al., 2025; Mankotia et al., 2023; W. Wang et al., 2021). Prior work from our group identified HY5 as an important player in iron homeostasis pathway, acting upstream of several key iron acquisition genes(Mankotia et al., 2023, 2024, 2025). To investigate whether the blue light mediated regulation of iron accumulation is dependent on HY5, we performed phenotypic analyses in both white and blue light conditions under +Fe and -Fe, using T-DNA insertional mutant of *HY5*. In WT seedlings, the iron deficiency led to increase in root length, an effect that was not observed in the *hy5* mutant, under both light conditions (Fig. 3a-b). Consistent with this, the Perl’s and Perls’ DAB staining revealed reduced Fe accumulation in *hy5* mutants compared to WT under both white and blue light (Fig. 3c). Moreover, quantitative Fe content analysis revealed that blue light induced increase in iron levels in WT seedlings, but *hy5* mutants accumulated significantly lower Fe levels in both light conditions (Fig. 3d). Phenotypic differences were also apparent in the shoot, where *hy5* mutant showed severe chlorosis under –Fe conditions specifically in blue light (Fig 3e). Quantification of total chlorophyll content revealed a significant reduction in *hy5* mutant compared to WT under both +Fe and –Fe conditions, with a more pronounced effect under blue light (Fig. 3e-f). A homolog of HY5, known as HYH have been shown to function redundantly with HY5 in the regulation of root growth, hypocotyl elongation and various aspects of light signaling. However, previous work has shown that HY5 regulates Fe homeostasis independent of HYH (Mankotia et al., 2023). This was further supported by Fe quantification analysis in *hyh* (WS) mutant. The blue light-induced increase in iron content observed in *hyh* (WS) was comparable to that of the wild type (WT, WS), whereas both *hy5* (WS) single and *hy5hyh* (WS) double mutants accumulated significantly less iron under both white and blue light conditions (Fig. S2).

**Fig. 3.**
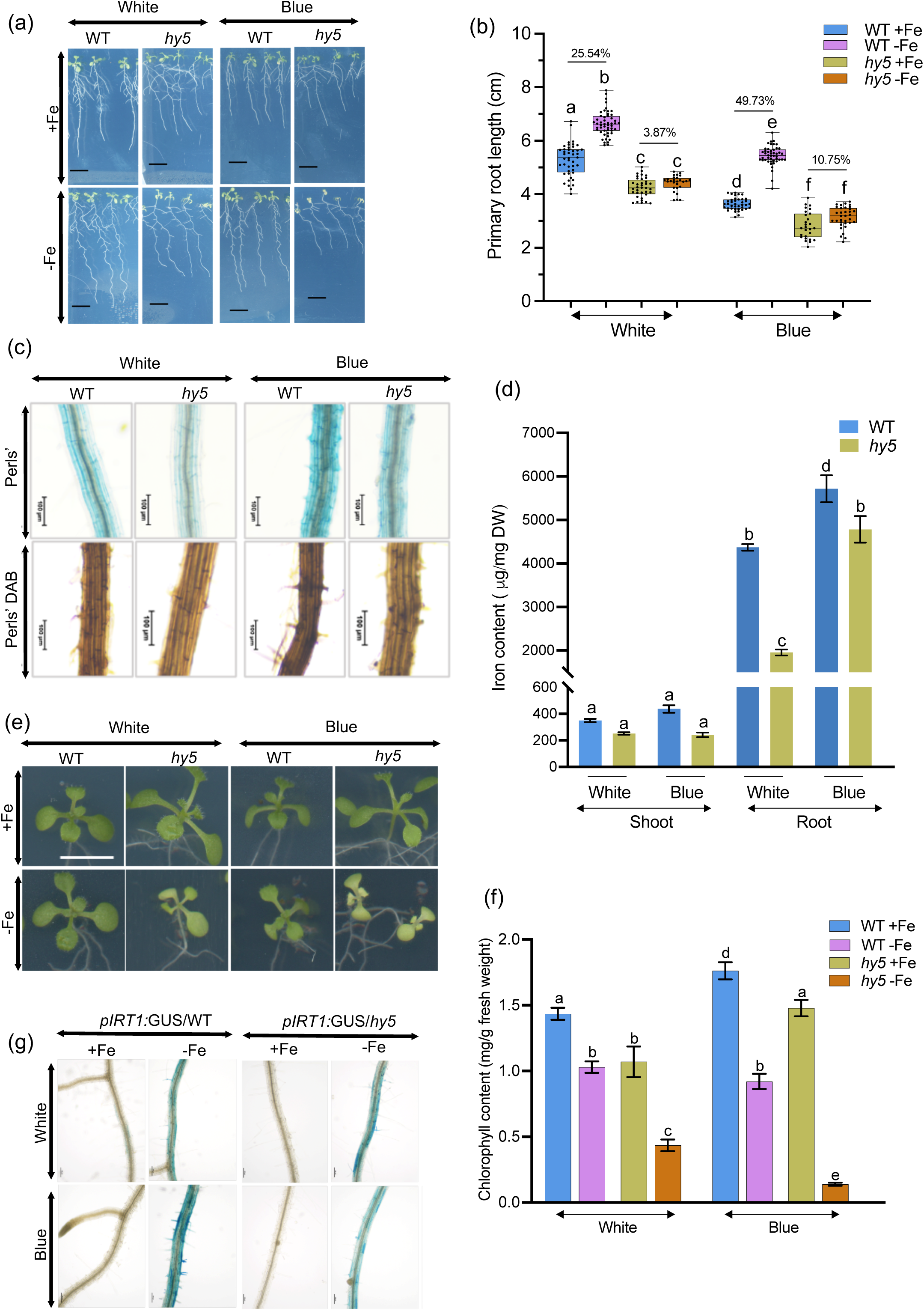
Blue light specific regulation of iron homeostasis is HY5 mediated. (a) Phenotype of WT and *hy5* plants grown on +Fe and -Fe media under white and blue light for 10 days. Scale bar: 1 cm. (b) Boxplot of root length of WT and *hy5* plants grown on +Fe and -Fe media under white and blue for 10 days. Means within each condition with the same letter are not significantly different according to one-way ANOVA followed by post hoc Tukey test, P < 0.05. using GraphPad PRISM 10. (c) Perls’ and Perls’ DAB stained maturation zone of WT and *hy5* plants grown on +Fe media under white and blue light for 5 days. Scale bar: 100 µM. (d) Iron content quantified spectrophotometrically in shoots and roots of WT and *hy5* plants grown on +Fe media under white and blue light for 15 days. The data represent average of three biological replicates, with three technical replicates of each. Each biological replicate consisted of ∼30 roots/shoots. Error bars represent ±SEM. DW-dry weight. Different letters (a, b, c, d) indicate significant differences, determined by one-way ANOVA followed by a post hoc Tukey Test (P≤0.05) using GraphPad PRISM 10. (e) Shoot phenotype of WT and *hy5* plants grown on +Fe and -Fe media under white and blue light for 10 days. Scale bar: 0.5 cm. (f) Total chlorophyll content in shoots of WT and *hy5* plants grown on +Fe and -Fe media under white and blue light for 10 days. Each biological replicate consisted of 5 shoots. Error bars represent ±SEM. Different letters (a, b, c, d) indicate significant differences, determined by one-way ANOVA followed by a post hoc Tukey Test (P≤0.05) using GraphPad PRISM 10. (g) Analysis of promoter activity in *pIRT1:GUS/WT* and *pIRT1:GUS/hy5* seedlings grown under white and blue light on +Fe, -Fe and transferred to –Fe (+300µM Fz) for 3 days. Scale bar: 100 µM.

To further investigate the molecular basis of this regulation, we checked the expression of *IRT1,* a key iron uptake gene, that is transcriptionally induced under Fe deficiency (Vert et al., 2002). We utilized *pIRT1:GUS* reporter lines in both WT and *hy5* mutant backgrounds and analyzed GUS activity in seedlings grown under white and blue light for 7 days on both +Fe and –Fe media. Under +Fe conditions GUS activity was undetectable in both genotypes and light conditions. In contrast, under

–Fe conditions, *pIRT1:GUS/WT* seedlings exhibited stronger GUS staining under blue light compared to white light, indicating enhanced *IRT1* induction. However, this induction was markedly reduced in *pIRT1:GUS/hy5* seedlings under both light conditions (Fig. 3g). These findings clearly demonstrate that HY5 is required for blue light mediated induction of *IRT1* expression under –Fe conditions, supporting the conclusion that blue light promotes iron accumulation and homeostasis via HY5-dependent regulatory pathways.

### *CRY1* is required for root growth under Fe deficiency and blue light induced Fe accumulation

Having established that blue light positively regulates Fe uptake in plants, we next sought to identify the photoreceptor(s) involved in mediating this response. Blue light receptors, cryptochromes (CRYs) and phototropins (PHOTs), are known to regulate various photomorphogenic responses (Lin, 2002). Cryptochromes are involved in regulation of circadian clock, cell elongation and flowering time, while phototropins on the other hand regulate phototropism, stomatal opening and chloroplast movement in response to different light intensities (Mawphlang & Kharshiing, 2017). To assess the role of these photoreceptors, we first examined the root elongation in cryptochrome and phototropin mutants under -Fe conditions. The *cry1* single mutant and *cry1cry2* double mutant failed to show typical increase in root length observed in WT seedlings under -Fe conditions. In contrast, *cry2* and *phot1phot2* mutants showed increase in root elongation comparable to the WT seedlings (Fig. 4a-b). Both the *cry1* and *cry1cry2* mutants showed severe reduction in Fe accumulation as compared to WT under both white and blue light (Fig. 4c). However, the iron content in *cry2* mutant was slightly lower than WT, while in *phot1phot2* mutant was similar to the WT and the blue light induced increase in Fe accumulation was also persistent in both *cry2* and *phot1phot2* mutants but not in *cry1* and *cry1cry2* (Fig. 4c). Spectrophotometric quantification of iron content in the shoots further confirmed that the both *cry1* and *cry1cry2* mutants accumulated less iron under both light conditions (Fig. 4d). These results suggest that CRY1 function is important for both root growth response under iron deficiency and blue light mediated Fe accumulation. However, a previous study in tomato demonstrated that phyB-mediated signaling plays a key role in regulating Fe uptake in tomato, revealed that *phyB-*mediated signaling plays a key role in regulating Fe uptake (Guo et al., 2021a). However, in Arabidopsis, the *phyB* mutant showed increased root length under Fe deficiency and Fe accumulation levels comparable to WT, supporting the notion that red light does not promote Fe accumulation and root elongation in response to Fe deficiency in Arabidopsis (Fig. S3).

**Fig. 4.**
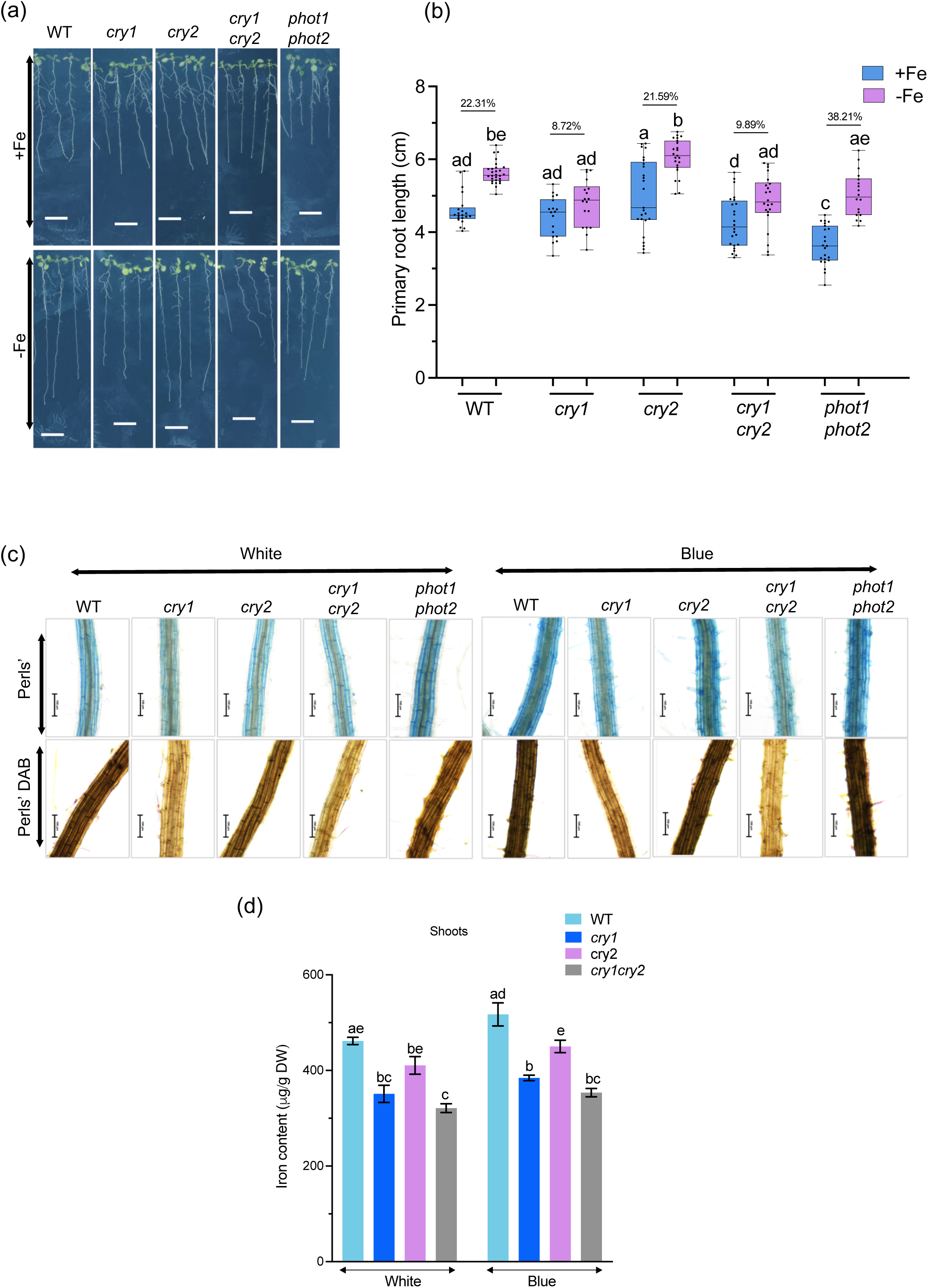
*CRY1* is required for blue light induced Fe accumulation and root growth under Fe deficiency. (a) Phenotype of WT, *cry1, cry2, cry1cry2* and *phot1phot2* seedlings grown on +Fe and - Fe media under white light for 10 days. Scale bar: 1 cm. (b) Boxplot of root length of WT, *cry1, cry2, cry1cry2* and *phot1phot2* seedlings grown on +Fe and -Fe media under white light for 10 days. Means within each condition with the same letter are not significantly different according to one-way ANOVA followed by post hoc Tukey test, P < 0.05 using GraphPad PRISM 10. (c) Perls’ and Perls’ DAB stained maturation zone of WT, *cry1, cry2, cry1cry2* and *phot1phot2* seedlings grown on +Fe media under white and blue light for 5 days. Scale bar: 100 µM. (d) Iron content quantified spectrophotometrically in shoots. Plants were grown on +Fe media under white and blue light for 15 days. The data represent average of three biological replicates, with three technical replicates of each. Each biological replicate consisted of ∼30 shoots. Error bars represent ±SEM. DW - dry weight. Different letters (a, b, c, d) indicate significant differences, determined by one-way ANOVA followed by a post hoc Tukey Test (P≤0.05) using GraphPad PRISM 10.

### Cryptochromes regulate the key Fe homeostasis genes

To understand the molecular basis cryptochromes mediated regulation of Fe uptake, we first examined the transcript levels of key Fe homeostasis genes. qRT-PCR analysis revealed that the expression of *IRT1, FRO2*, and *HY5* was markedly reduced in *cry1*, *cry2*, and *cry1 cry2* mutants under iron-deficient conditions compared to WT under white (Fig. 5a), being more prominent in blue light (Fig 5b). Interestingly, HY5 expression was slightly induced under white light in *cry1* single mutant but this induction was not seen under blue light (Fig. 5a-b). Furthermore, the *cry2* mutant exhibited transcript levels similar to those of the *cry1cry2* double mutant, suggesting a predominant role of CRY1 in regulating Fe uptake and root elongation, while CRY2 may contribute primarily at the transcriptional level within this pathway. To further assess gene expression *in-vivo,* we analyzed *pIRT1:GUS* transgenic lines in *cry1cry2* mutant background. In *pIRT1:GUS* seedlings, GUS activity was substantially reduced in *cry1cry2* mutants under –Fe conditions, and the blue light–induced enhancement of *IRT1* expression was also absent (Fig. 5c). Additionally, we assessed IRT1 protein level under +Fe and -Fe conditions in WT and *cry1cry2* mutants. Immunoblot results showed that IRT1 protein levels were less strongly induced in *cry1cry2* under -Fe in white light (Fig. 5d). Next, we measured the ferric chelate reductase (FRO2) activity and found that *cry1* exhibited FRO2 activity similar to WT, whereas both *cry2* and *cry1cry2* mutants showed significantly reduced activity, particularly under blue light (Fig. 5e). Consistently, confocal imaging of *pFRO2:FRO2:mCherry* revealed that FRO2 accumulation was not upregulated under –Fe conditions in in *cry1cry2* mutant background (Fig. S4). Taken together, these results suggest that CRY1/CRY2, in concert with HY5, forms a blue light–responsive regulatory module that positively regulates Fe uptake and homeostasis in *Arabidopsis*. While CRY1 primarily mediates physiological responses, CRY2 is likely to fine-tune transcriptional regulation of Fe uptake genes.

**Fig. 5.**
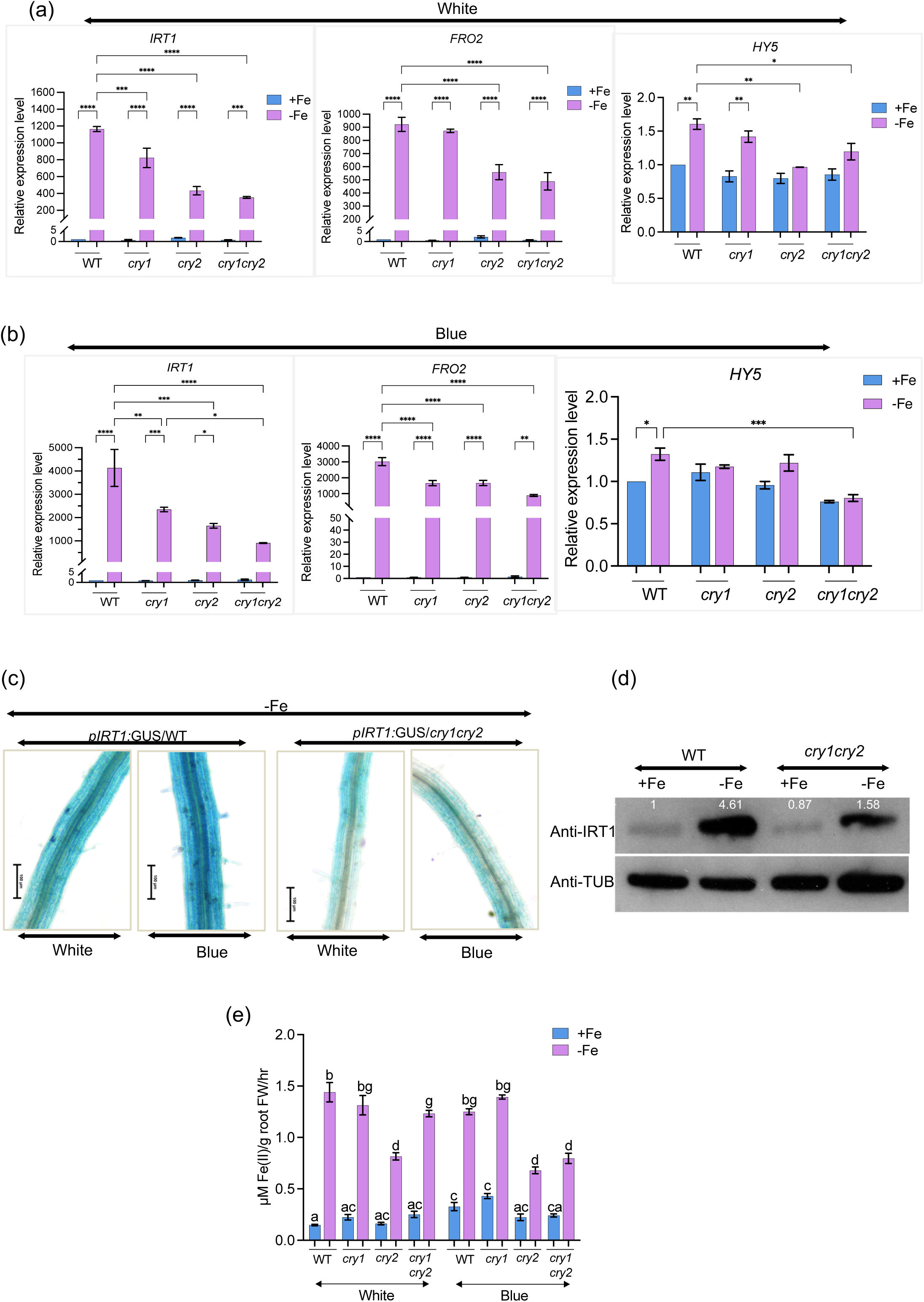
Cryptochromes regulate the key Fe homeostasis genes. (a) Expression levels of *IRT1*, *FRO2 and HY5 .* Plants were grown under white light for 5 days on +Fe and then transferred to +Fe and –Fe(+300 µM Fz) for 3 days. WT used as control for each treatment. * represents significant difference according to one-way ANOVA followed by post hoc Tukey test, P < 0.05. using GraphPad PRISM 10. (b) Expression levels of *IRT1*, *FRO2 and HY5.* Plants were grown under blue light for 5 days on +Fe and then transferred to +Fe and –Fe(+300 µM Fz) for 3 days under white light. WT used as control for each treatment. * represents significant difference according to one-way ANOVA followed by post hoc Tukey test, P < 0.05. using GraphPad PRISM 10. (c) Analysis of promoter activity in *pIRT1:GUS/WT* and *pIRT1:GUS* crossed with *cry1cry2.* Seedlings were grown under white and blue light on -Fe. Scale bar: 100 µM. (d) Western Blot showing IRT1 protein levels in WT and *cry1cry2* seedlings grown on +Fe media for five days and then transferred to +Fe and –Fe (+ 300 µM Fz) media for 3 days, under white light. Tubulin - Internal Control. The values above each band depicts the intensities after normalization with internal control, followed by normalization using WT +Fe as a control. (e) FRO2 activity in WT, *cry1, cry2* and *cry1cry2* roots grown for five days on +Fe media and then transferred to +Fe and -Fe (+300μM Fz) media for 3 days, under white and blue light. Means with the same letter are not significantly different according to one-way ANOVA followed by post hoc Tukey test, P < 0.05 using GraphPad PRISM 10. Error bars represent ±SEM.

### CRY–HY5 signaling coordinates blue light regulation of iron homeostasis

To further dissect the genetic interaction between CRY1, CRY2 and HY5 in regulating Fe homeostasis under blue light, we generated *hy5cry1*, *hy5cry2*, and *hy5cry1cry2* double and triple mutants. The *hy5cry1cry2* triple mutant did not show the iron deficiency-mediated increase in root length (Fig. 6a-b). The iron accumulation in the triple mutant was also markedly reduced and the blue light mediated increase was absent, similar to *hy5* and *cry1cry2* mutants (Fig. 6c). To further confirm these findings and understand the shoot specific role of CRYs and HY5, we performed reciprocal grafting experiments using WT*, cry1cry2,* and *hy5* plants. We found that, across all grafting combinations, plants with *cry1cry2* or *hy5* shoots accumulated less iron (Fig. 6d). Notably, grafting WT roots to *cry1cry2* or *hy5* shoots did not restore Fe levels similar to WT plants, indicating that blue light perception occurs primarily in the shoot and that CRY1-CRY2 and HY5 function are required for the signaling cascade. Chlorophyll measurements further supported these findings (Fig. S5e, S6e). The Fe deficiency-induced root elongation response was not observed in *hy5cry1* and *hy5cry1cry2* mutants, consistent with the phenotypes of the corresponding single and double mutants under both light conditions (Fig. S5a–b, S6a–b). Similarly, the iron accumulation was also affected in the single and double mutants (Fig. S5c, S6c), however, the *hy5cry2* double mutant showed a mild increase under blue light (Fig. S6c). Double and triple mutants showed a more pronounced reduction in total chlorophyll content compared to single mutants under both white (Fig. S5d-e) and blue light (Fig. S6d-e). Altogether, our work has shown that light perception via CRY–HY5 signaling is crucial for blue light–mediated iron homeostasis, as *hy5cry1cry2* mutants showed impaired molecular and physiological effects.

**Fig. 6.**
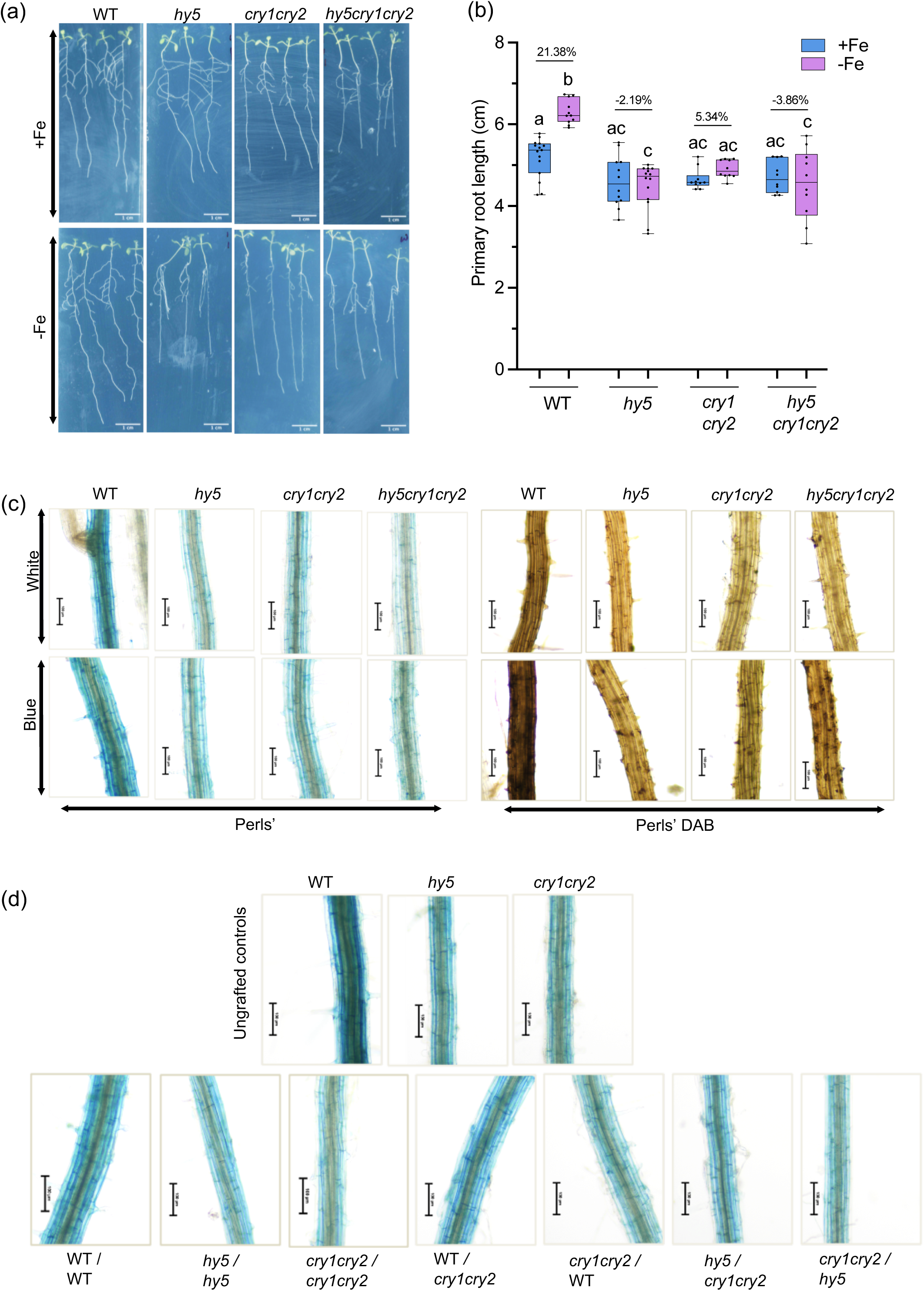
CRY–HY5 signaling coordinates blue light regulation of iron homeostasis. (a) Phenotype of WT, *hy5, cry1cry2 and hy5cry1cry2* plants grown on +Fe media for 5 days and then transferred to +Fe and -Fe media under white and blue light for 5 days. The *hy5cry1cry2* triple mutants were screened on basis of hypocotyl length. Scale bar: 1 cm. (b) Boxplot of root length of WT, *hy5, cry1cry2 and hy5cry1cry2* plants grown on +Fe media for 5 days and then transferred to +Fe and -Fe media under white and blue light for 5 days. Means within each condition with the same letter are not significantly different according to one-way ANOVA followed by post hoc Tukey test, P < 0.05. using GraphPad PRISM 10. (c) Perls’ and Perls’ DAB stained maturation zone of WT, *hy5, cry1cry2 and hy5cry1cry2* plants grown on +Fe media under white and blue light for 5 days. The *hy5cry1cry2* triple mutants were screened on basis of hypocotyl length. Scale bar: 100 µM. (d) Perls’ stained maturation zone of different grafting combinations. 7 day old WT, *hy5* and *cry1cry2* seedlings grown on +Fe media were used for grafting. All the grafting combinations were allowed to grow for another 5 days and successful grafts were used for Perls’ staining. Scale bar: 100 µM.

## Discussion

Plants being sessile in nature depend on a fine tuning of external environmental changes with internal molecular cues to regulate various aspects of their growth and development. Among these cues, light is one of the most influential factors, orchestrating diverse physiological processes through shoot-localized perception by different photoreceptors and systemic signaling. While light’s role in shoot morphogenesis is well established, growing evidence reveals that light perception in the shoot also mediates long-distance regulatory effects on root development and nutrient homeostasis (Lee et al., 2016; Y. Zhang et al., 2019). These systemic effects are often mediated by mobile transcription factors, such as HY5, which translocate from the shoot to the root and modulate gene expression in response to light. Light regulates HY5 dependent expression of multiple genes involved in stress, metabolism and phytohormone signaling (Zhang et al., 2019).

In addition to its well established roles in photomorphogenesis, thermotolerance, and nutrient signaling, HY5 has been shown to mediate light-dependent root responses to phosphorus and nitrogen availability (Chen et al., 2016; Gao et al., 2021b; Yeh et al., 2020)Our previous work demonstrated that HY5 positively regulates iron homeostasis in Arabidopsis by directly targeting key iron acquisition genes (Mankotia et al., 2023). However, the light-dependent regulation of iron uptake and the upstream signaling mechanisms remained poorly understood.

In this study, we uncover a critical role for blue light in regulating Fe uptake and homeostasis via a CRY–HY5 signaling module. We show that blue light, but not red light, promotes Fe accumulation in Arabidopsis and this response is tightly coupled to the shoot’s exposure to light, reflecting the natural growth conditions where only aerial tissues are illuminated (Fig. 1–2). Under Fe-deficient conditions, blue light significantly enhances HY5 protein accumulation (Fig. 2f), indicating that light quality modulates HY5 stability or expression under Fe deficiency.

Our phenotypic and molecular analyses of *hy5* mutants further confirmed that HY5 function is essential for the blue light-mediated increase in Fe content and root elongation under Fe-deficient conditions (Fig. 3). To identify the upstream photoreceptors involved in this response, we examined mutants of multiple photoreceptors. Interestingly, *cry1* and *cry1cry2* mutants displayed strong defects in Fe accumulation, while *cry2* mutants showed only mild reductions and *phot1phot2* and *phyB* mutants behaved similarly to WT (Fig. 4, S3). These findings suggest that CRY1 plays a predominant role in blue light-mediated Fe accumulation, with CRY2 contributing a modulatory function.

Interestingly, while *cry1* mutants showed normal FRO2 enzymatic activity, *cry2* and *cry1cry2* mutants exhibited reduced activity (Fig. 5e), suggesting to a functional divergence between CRY1 and CRY2. This was further supported by expression analysis of key Fe homeostasis genes (*IRT1, FRO2*, and *HY5*), where transcript levels were strongly reduced in *cry2* and *cry1cry2* backgrounds under both white and blue light (Fig. 5a-b). However, under blue light, the difference becomes much more significant, where *cry1, cry2, cry1cry2* showed reduced transcript levels of *IRT1, FRO2*, and *HY5* (Fig. 5b). Reporter lines also confirmed significantly decreased expression of *FRO2* and *IRT1* in cryptochrome mutants, particularly under blue light (Fig. 5c, S4). Moreover, immunoblot analysis showed that IRT1 protein accumulation was less induced in *cry1cry2* under -Fe in white light (Fig. 5d). Together, these results indicate that both CRY1 and CRY2 are necessary for optimum activation of the Fe uptake machinery.

Genetic analysis *of hy5cry1, hy5cry2, and hy5cry1cry2* mutants further supports the conclusion that CRY1 and CRY2 act upstream of HY5 in this pathway. All double and triple mutants showed reduced Fe accumulation, lower chlorophyll content, and failed to elongate roots in response to Fe deficiency under both white and blue light (Fig. 6, S6, S7). Consistently, grafting experiments further revealed that the blue light perception occurs primarily in the shoot, as *cry1cry2* shoots were unable to restore Fe accumulation even when grafted onto WT roots (Fig. 6d). These findings strongly support a model where light perception in the shoot via cryptochromes triggers HY5-mediated long-distance signaling to regulate root Fe acquisition.

**Fig. 7.**
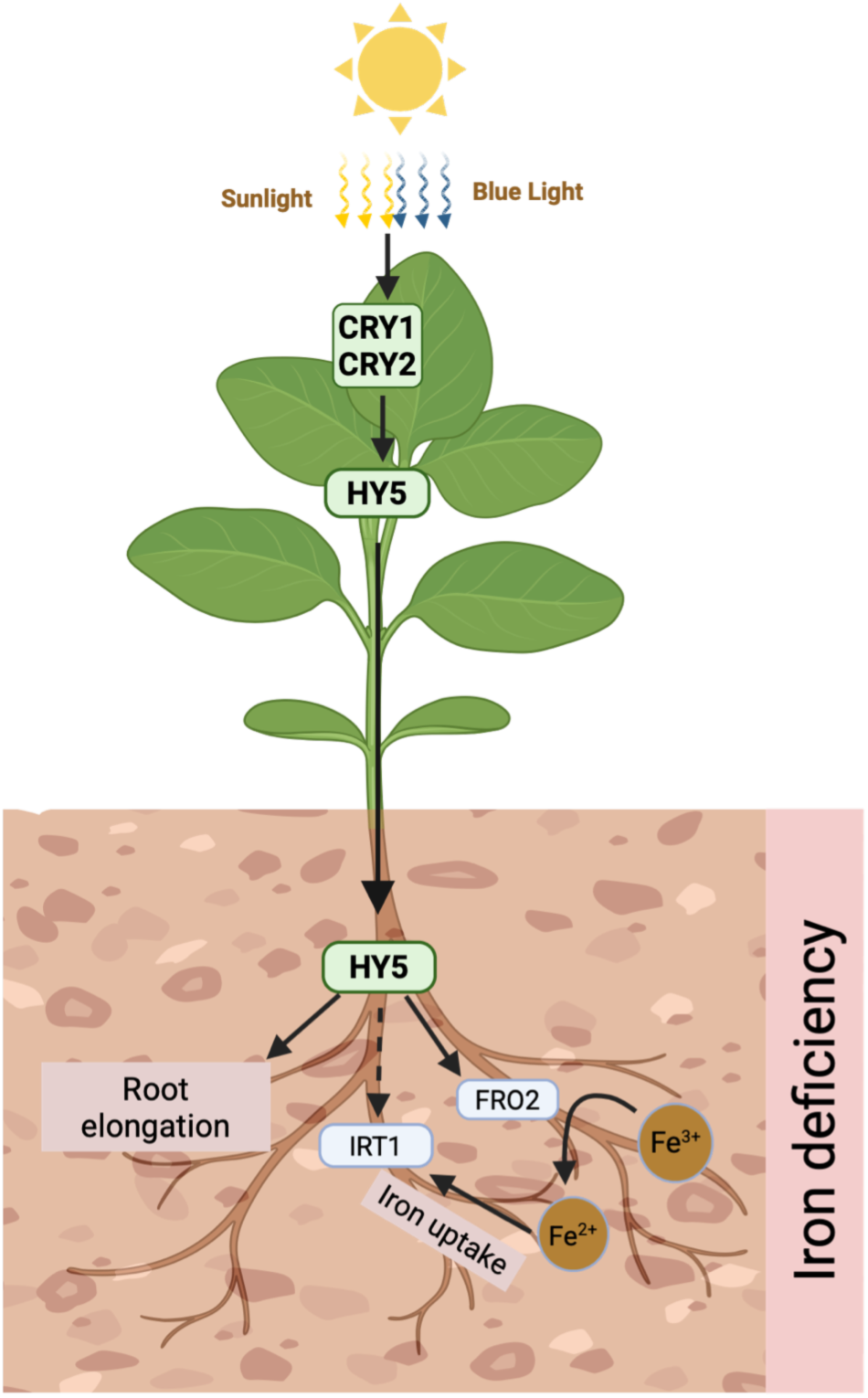
Blue light mediates iron homeostasis via CRY-HY5 module. Blue light perceived by CRY1 and CRY2 activates HY5 in the shoots, which then migrates to roots. In roots, HY5 regulates iron homeostasis by positively regulating *FRO2* directly and *IRT1* expression indirectly, which leads to increased iron uptake and accumulation, thus regulating iron homeostasis in Arabidopsis. Figure created using BioRender.

Taken together, our results establish a mechanistic framework in which blue light perceived by CRY1 and CRY2 in the shoot enhances HY5 accumulation, which subsequently activates downstream Fe uptake genes in the root (Fig. 7). This CRY–HY5 module integrates light signals with nutrient status, enabling plants to fine-tune Fe acquisition under fluctuating environmental conditions.

Beyond its mechanistic insights, this work offers translational potential in the context of controlled-environment agriculture. As indoor and hydroponic farming systems become increasingly important for sustainable food production, optimizing light signals (specifically blue light exposure) could be a promising strategy to enhance Fe content and improve the nutritional quality of crops. Supporting this, our preliminary experiments in mung bean (*Vigna radiata*) demonstrated enhanced Fe accumulation under blue light (Fig. S7), suggesting that similar mechanism may operate in other species.

In summary, we demonstrate that blue light plays a crucial role in regulating Fe homeostasis through a CRY1/CRY2–HY5 signaling axis. This study advances our understanding of how aboveground light signals coordinate belowground nutrient acquisition and opens new avenues for light-based strategies to enhance micronutrient uptake in crops.

## Acknowledgement

We thank Chentao Lin for providing *cry1, cry2, cry1cry2* seeds. We are also thankful to Rumen Ivanov and Petra Bauer for *pIRT1:GUS* seeds. We are also grateful to Roman Ulm for providing WS, *hyh, hy5, hyh/ hy5, cry1cry2, phot1phot2* seeds. We also thank Ferenc Nagy, Maria Agustina Mazzella, Sourav Datta and Sreeramaiah Gangappa for *phyB-9* seeds. We further extend our thanks to On Sun Lau for providing *pHY5:HY5:YFP/hy5* seeds. We are thankful to all the members of SBS lab. SBS acknowledges intramural funding support from Indian Institute of Science Education and Research (IISER) Mohali. SBS acknowledges the support through grant no. BT/PR51324/AGIII/103/1480/2023 from the Department of Biotechnology (DBT). SBS also acknowledges the Science and Engineering Research Board (SERB) for research funding (CRG/223 2022/003773). The SBS laboratory is also supported by the Indo-French Centre for the Promotion of Advanced Research (IFCPAR/ CEFIPRA) under project 68T06-1. PJ acknowledge PhD fellowship received from Council for Scientific and Industrial Research (CSIR), Govt. of India.

## Competing Interests

The authors declare no competing interests.

## Author contributions

Pooja Jakhar: Conceptualization; Data curation; Formal analysis; Validation; Investigation; Visualization; Methodology; Writing—original draft, review and editing. Samriti Mankotia: Conceptualization, Writing—review and editing. Shubham Ladhwal: Data curation; Formal analysis; Writing—review. Nisha Bodh: Methodology; Data curation. Krishna Dhanawat: Methodology; Data curation. Ajay K. Pandey: Formal analysis, Writing—review and editing.

Santosh B. Satbhai: Conceptualization; Formal analysis; Supervision; Funding acquisition; Investigation; Methodology; Project administration; Writing—original draft, review and editing.

**Fig. S1 Blue light positively regulates iron accumulation in Arabidopsis.**

Perls’ and Perls’ DAB stained maturation zone of WT seedlings grown on +Fe media under white light for 5 days and then transferred to blue and red light for 24 hours, 48 hours and 72 hours. Scale bar: 100 µM.

**Fig. S2 Blue light mediated iron accumulation is via HY5, independent of HYH.**

Perls’ and Perls’ DAB stained maturation zone of wild type (WS), *hy5* (WS), *hyh* (WS) and *hy5hyh* (WS) seedlings grown on +Fe media under white light for 5 days. Scale bar: 100 µM.

**Fig. S3 Phytochrome B is not involved in iron homeostasis.**

(a) Phenotype of WT and *phyB* seedlings grown on +Fe and -Fe media under white light for 10 days. Scale bar:1 cm. (b) Boxplot of root length of WT and *phyB* seedlings grown on +Fe and -Fe media under white light for 10 days. Means within each condition with the same letter are not significantly different according to one-way ANOVA followed by post hoc Tukey test, P < 0.05. using GraphPad PRISM 10. (c) Perls’ and Perls’ DAB stained maturation zone of WT and *phyB* seedlings grown on +Fe media under white light for 5 days. Scale bar: 100 µM.

**Fig. S4 Cryptochromes are required for iron deficiency induced expression of *FRO2*.**

(a) Confocal images of *pFRO2:FRO2:mCherry/frd1* seedlings and cross of *cry1cry2* with *pFRO2:FRO2:mCherry/frd1.* Seedlings were grown on +Fe media for five days and then transferred to +Fe and –Fe (+300 µM Fz) media for 3 days. A representative image from maturation zone is shown for each. Scale bar: 100µm. Autofluorescence is detected in the near-infrared (NIR) range (650-700 nm) when excited by a blue laser. (b) Quantification of mean intensity of mCherry signal in (a). Error bars represent SEM. Different alphabets indicate significant difference according to one-way ANOVA followed by post hoc Tukey test, P < 0.05 using GraphPad PRISM 10.

**Fig. S5 Cryptochrome controls light mediated regulation of iron homeostasis via HY5.**

(a) Phenotype of WT, *hy5, cry1, cry2, cry1cry2, hy5cry1, hy5cry* and *hy5cry1cry2* plants grown on +Fe media for 5 days and then transferred to +Fe and -Fe media under white light for 5 days. The double and triple mutants were screened on basis of hypocotyl length. Scale bar:1 cm. (b) Boxplot of root length of WT, *hy5, cry1, cry2, cry1cry2, hy5cry1, hy5cry* and *hy5cry1cry2* plants grown on +Fe media for 5 days and then transferred to +Fe and -Fe media under white light for 5 days. Means within each condition with the same letter are not significantly different according to one-way ANOVA followed by post hoc Tukey test, P < 0.05. using GraphPad PRISM 10. (c) Perls’ and Perls’ DAB stained maturation zone of WT, *hy5, cry1, cry2, cry1cry2, hy5cry1, hy5cry* and *hy5cry1cry2* plants grown on +Fe media under white light for 5 days. The double and triple mutants were screened on basis of hypocotyl length. Scale bar: 100 µM. (d) Shoot phenotype of WT, *hy5, cry1, cry2, cry1cry2, hy5cry1, hy5cry* and *hy5cry1cry2* plants grown on +Fe and -Fe media under white for 10 days. Scale bar: 0.5 cm. (e) Percentage reduction in chlorophyll in shoots of WT, *hy5, cry1, cry2, cry1cry2, hy5cry1, hy5cry2* and *hy5cry1cry2* plants grown on +Fe and -Fe media under white light for 10 days. Each biological replicate consisted of 5 shoots. Error bars represent ±SEM. Different letters (a, b, c, d) indicate significant differences, determined by one-way ANOVA followed by a post hoc Tukey Test (P≤0.05) using GraphPad PRISM 10.

**Fig. S6 CRY1 seems to be main blue light photoreceptor upstream to HY5 in regulating iron uptake and root growth** (a) Phenotype of WT, *hy5, cry1, cry2, cry1cry2, hy5cry1, hy5cry* and *hy5cry1cry2* plants grown on +Fe media for 5 days and then transferred to +Fe and -Fe media under blue light for 5 days. The double and triple mutants were screened on basis of hypocotyl length. Scale bar: 1 cm. (b) Boxplot of root length of WT, *hy5, cry1, cry2, cry1cry2, hy5cry1, hy5cry* and *hy5cry1cry2* plants grown on +Fe media for 5 days and then transferred to +Fe and -Fe media under blue light for 5 days. Means within each condition with the same letter are not significantly different according to one-way ANOVA followed by post hoc Tukey test, P < 0.05. using GraphPad PRISM 10. (c) Perls’ and Perls’ DAB stained maturation zone of WT, *hy5, cry1, cry2, cry1cry2, hy5cry1, hy5cry* and *hy5cry1cry2* plants grown on +Fe media under blue light for 5 days. The double and triple mutants were screened on basis of hypocotyl length. Scale bar: 100 µM. (d) Shoot phenotype of WT, *hy5, cry1, cry2, cry1cry2, hy5cry1, hy5cry* and *hy5cry1cry2* plants grown on +Fe and -Fe media under blue for 10 days. Scale bar: 0.5 cm. (e) Percentage reduction in chlorophyll in shoots of WT, *hy5, cry1, cry2, cry1cry2, hy5cry1, hy5cry* and *hy5cry1cry2* plants grown on +Fe and -Fe media under blue light for 10 days. Each biological replicate consisted of 5 shoots. Error bars represent ±SEM. Different letters (a, b, c, d) indicate significant differences, determined by one-way ANOVA followed by a post hoc Tukey Test (P≤0.05) using GraphPad PRISM 10.

**Fig. S7 Moong bean sprouted in dark for 2 days were transferred to liquid Hoagland media for 5 days under different lights.** (a) Perls’ stained seedlings of Mung bean, sprouted in dark for 2 days and then transferred to Fe-sufficient liquid Hoagland media under different lights for 5 days. Scale bar: 1 cm. (b) Perls’ stained maturation zone of Mung bean marked in (a) as viewed under microscope. Scale bar: 100 µM.

## Notes

### Competing Interest Statement

The authors have declared no competing interest.

