## Supporting information for "Blue Light Promotes Root Iron Acquisition via a Shoot CRY–HY5 Signaling Module in Arabidopsis"

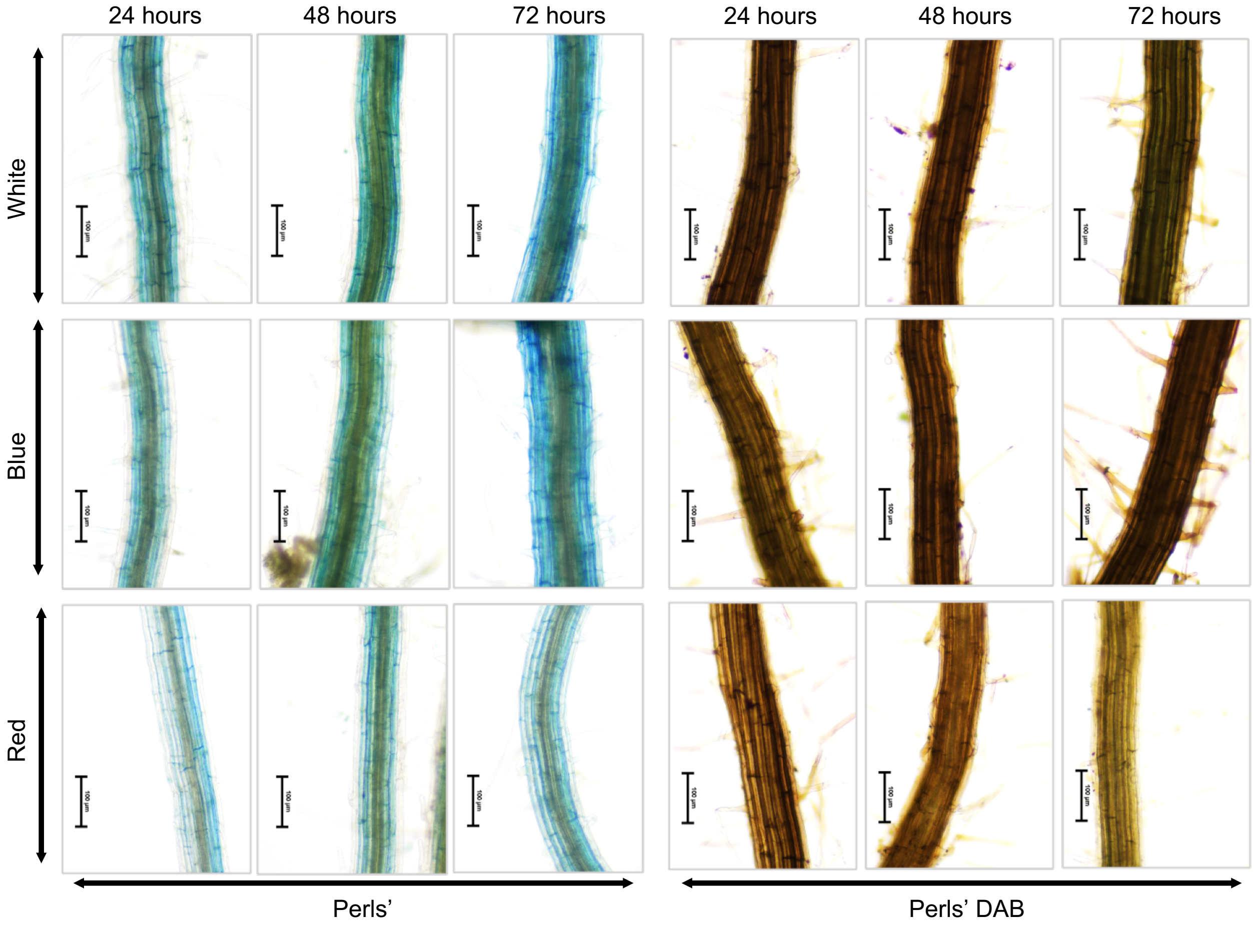


**Fig. S1 Blue light positively regulates iron homeostasis in Arabidopsis.**
Perls' and Perls' DAB stained maturation zone of WT seedlings grown on +Fe media under white light for 5 days and then transferred to blue and red light for 24 hours, 48 hours and 72 hours. Scale bar: 100 μM.

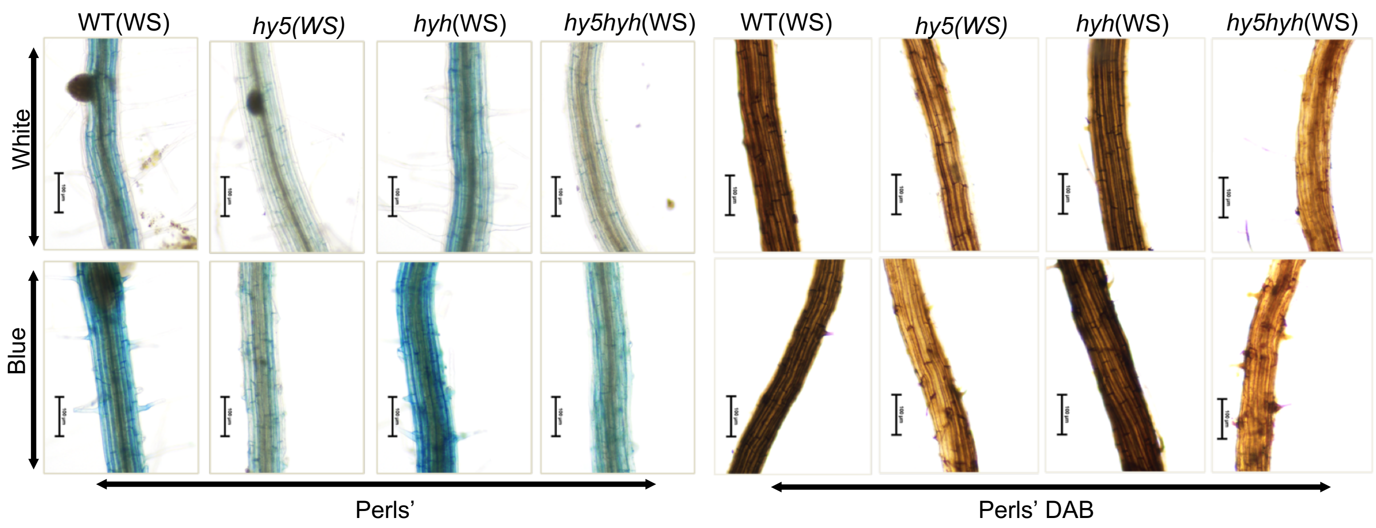


**Fig. S2 Blue light mediated iron accumulation is via HY5, independent of HYH.**
Perls' and Perls' DAB stained maturation zone of wild type (WS), *hy5(*WS), *hyh*(WS) and *hy5hyh*(WS) seedlings grown on +Fe media under white light for 5 days. Scale bar: 100 μM.

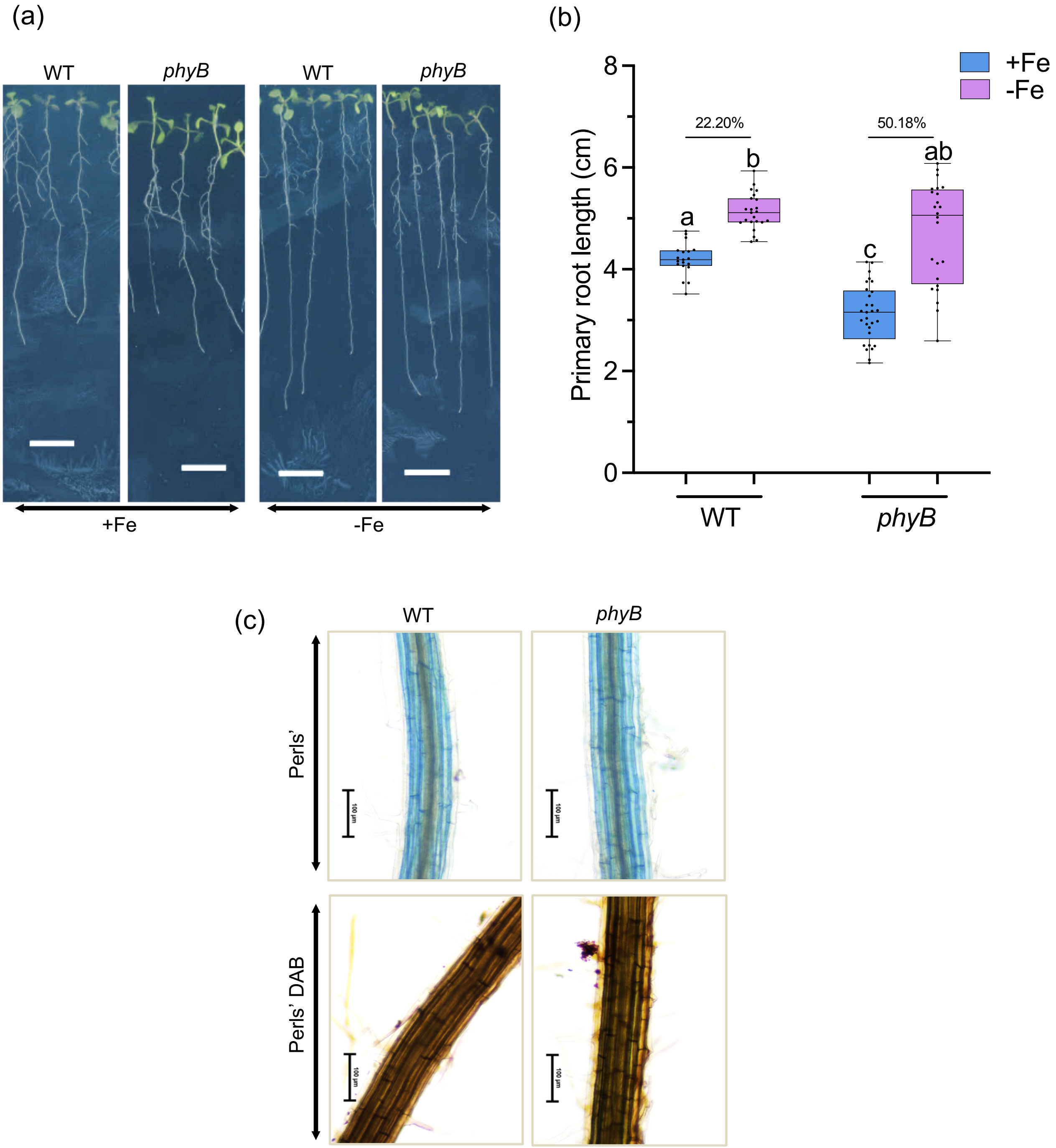


**Fig. S3 Phytochrome B is not involved in iron homeostasis.**
(a) Phenotype of WT and *phyB* seedlings grown on +Fe and -Fe media under white light for 10 days. Scale bar: 1 cm. (b) Boxplot of root length of WT and *phyB* seedlings grown on +Fe and -Fe media under white light for 10 days. Means within each condition with the same letter are not significantly different according to one-way ANOVA followed by post hoc Tukey test, $P < 0.05$. using GraphPad PRISM 10. (c) Perls' and Perls' DAB stained maturation zone of WT and *phyB* seedlings grown on +Fe media under white light for 5 days. Scale bar: 100 μM.

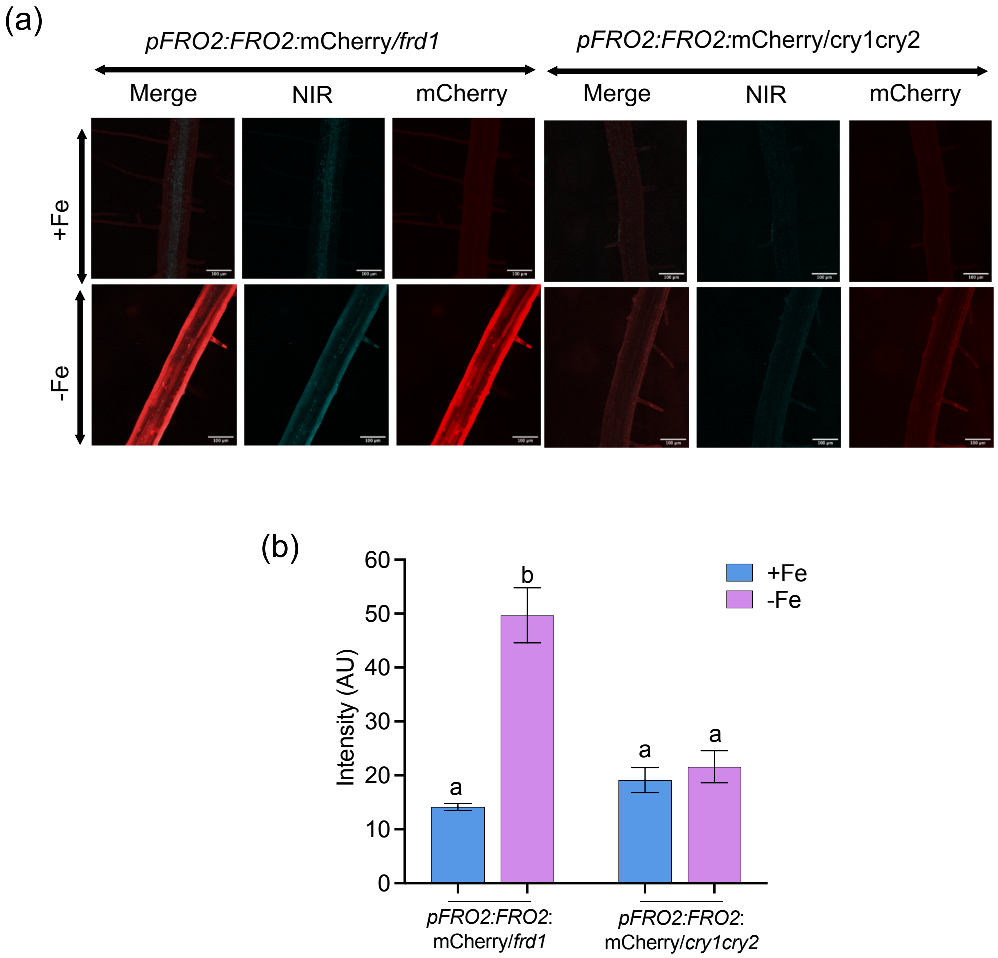


**Fig. S4 Cryptochromes are required for iron deficiency induced expression of *FRO2*.**
(a) Confocal images of *pFRO2:FRO2:mCherry/frd1* seedlings and cross of *cry1cry2* with *pFRO2:FRO2:mCherry/frd1*. Seedlings were grown on +Fe media for five days and then transferred to +Fe and –Fe (+300 μM Fz) media for 3 days. A representative image from maturation zone is shown for each. Scale bar: 100μm. Autofluorescence is detected in the near-infrared (NIR) range (650-700 nm) when excited by a blue laser. (b) Quantification of mean intensity of mCherry signal in (a). Error bars represent SEM. Different alphabets indicate significant difference according to one-way ANOVA followed by post hoc Tukey test, $P < 0.05$ using GraphPad PRISM 10.

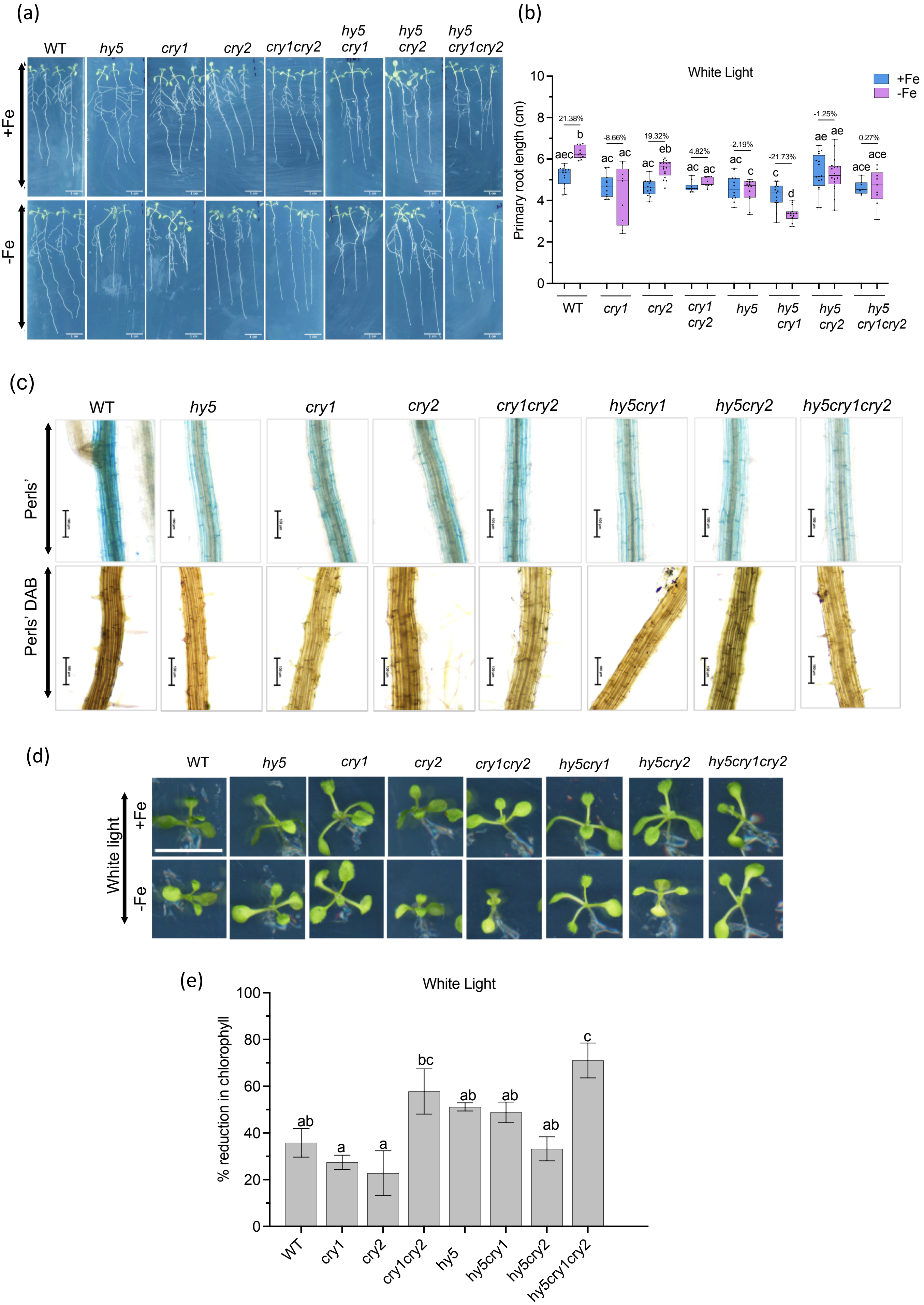
(a)
WT
hy5
cry1
cry2
cry1cry2
hy5 cry1
hy5 cry2
hy5 cry1cry2
+Fe
-Fe
(b)
White Light
+Fe
-Fe
Primary root length (cm)
10
8
6
4
2
0
21.38%
-8.66%
19.32%
4.82%
-2.19%
-21.73%
-1.25%
0.27%
aec
b
ac
ac
ac
eb
ac
ac
ac
c
c
d
ae
ae
ace
ace
WT
cry1
cry2
cry1 cry2
hy5
hy5 cry1
hy5 cry2
hy5 cry1cry2
(c)
WT
hy5
cry1
cry2
cry1cry2
hy5cry1
hy5cry2
hy5cry1cry2
Perls'
Perls' DAB
(d)
WT
hy5
cry1
cry2
cry1cry2
hy5cry1
hy5cry2
hy5cry1cry2
White light
+Fe
-Fe
(e)
White Light
% reduction in chlorophyll
100
80
60
40
20
0
ab
a
a
bc
ab
ab
ab
c
WT
cry1
cry2
cry1cry2
hy5
hy5cry1
hy5cry2
hy5cry1cry2

**Fig. S5 Cryptochrome controls light mediated regulation of iron homeostasis via HY5.**
(a) Phenotype of WT, *hy5, cry1, cry2, cry1cry2, hy5cry1,hy5cry* and *hy5cry1cry2* plants grown on +Fe media for 5 days and then transferred to +Fe and -Fe media under white light for 5 days. The double and triple mutants were screened on basis of hypocotyl length. Scale bar: 1 cm. (b) Boxplot of root length of WT, *hy5, cry1, cry2, cry1cry2, hy5cry1,hy5cry* and *hy5cry1cry2* plants grown on +Fe media for 5 days and then transferred to +Fe and -Fe media under white light for 5 days. Means within each condition with the same letter are not significantly different according to one-way ANOVA followed by post hoc Tukey test, $P < 0.05$. using GraphPad PRISM 10. (c) Perls' and Perls' DAB stained maturation zone of WT, *hy5, cry1, cry2, cry1cry2, hy5cry1,hy5cry* and *hy5cry1cry2* plants grown on +Fe media under white light for 5 days. The double and triple mutants were screened on basis of hypocotyl length. Scale bar: 100 μM. (d) Shoot phenotype of WT, *hy5, cry1, cry2, cry1cry2, hy5cry1,hy5cry* and *hy5cry1cry2* plants grown on +Fe and -Fe media under white for 10 days. Scale bar: 0.5 cm. (e) Percentage reduction in chlorophyll in shoots of WT, *hy5, cry1, cry2, cry1cry2, hy5cry1,hy5cry* and *hy5cry1cry2* plants grown on +Fe and -Fe media under white light for 10 days. Each biological replicate consisted of 5 shoots. Error bars represent ±SEM. Different letters (a, b, c, d) indicate significant differences, determined by one-way ANOVA followed by a post hoc Tukey Test ($P \leq 0.05$) using GraphPad PRISM 10.

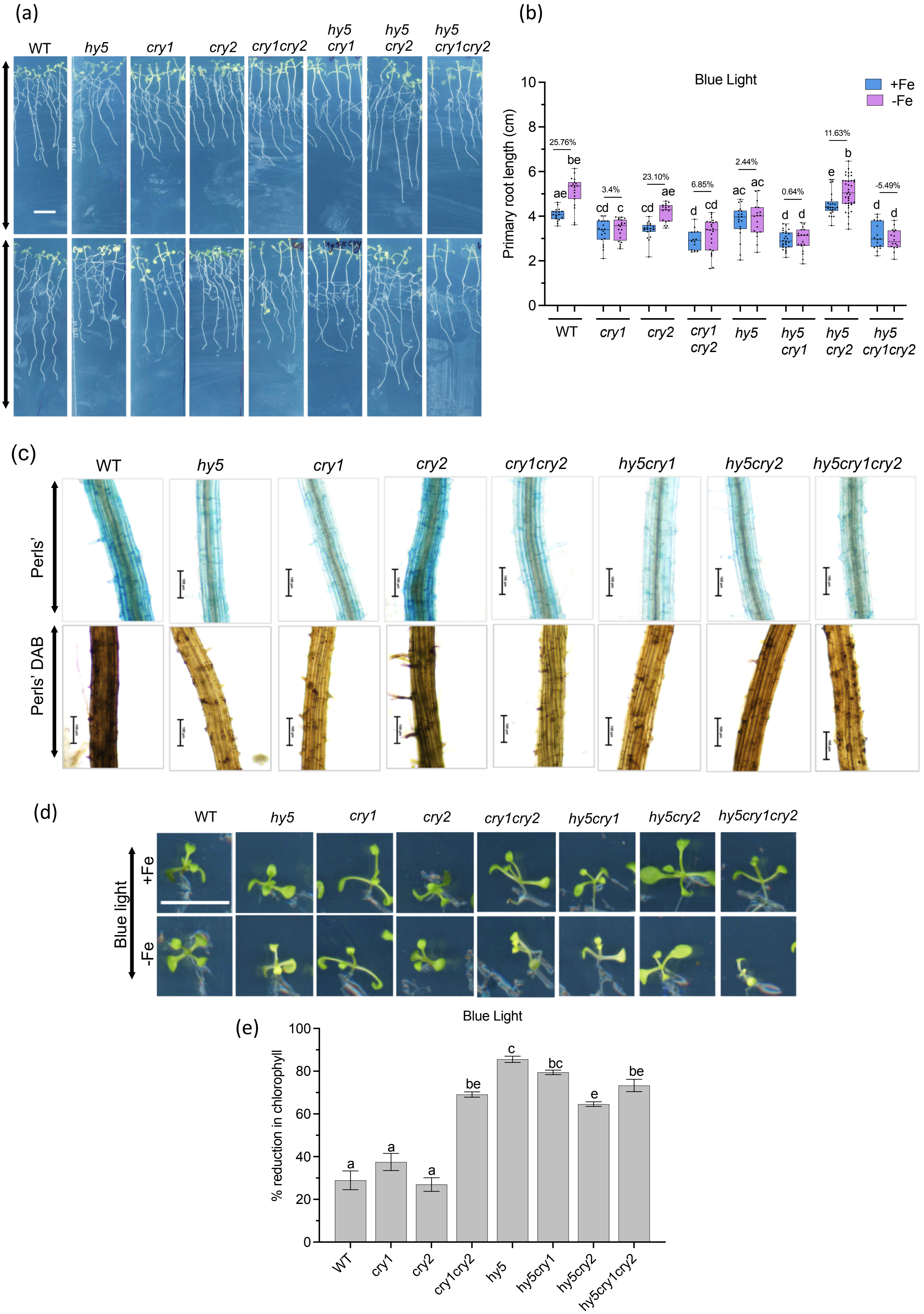

(a)
WT
hy5
cry1
cry2
cry1cry2
hy5 cry1
hy5 cry2
hy5 cry1cry2
+Fe
-Fe
(b)
Blue Light
+Fe
-Fe
Primary root length (cm)
10
8
6
4
2
0
25.76%
3.4%
23.10%
6.85%
2.44%
0.64%
11.63%
-5.49%
ae
be
cd
c
cd
ae
d
cd
ac
ac
d
d
e
b
d
d
WT
cry1
cry2
cry1 cry2
hy5
hy5 cry1
hy5 cry2
hy5 cry1cry2
(c)
WT
hy5
cry1
cry2
cry1cry2
hy5cry1
hy5cry2
hy5cry1cry2
Perls'
Perls' DAB
(d)
WT
hy5
cry1
cry2
cry1cry2
hy5cry1
hy5cry2
hy5cry1cry2
Blue light
+Fe
-Fe
(e)
Blue Light
% reduction in chlorophyll
100
80
60
40
20
0
a
a
a
be
c
bc
e
be
WT
cry1
cry2
cry1cry2
hy5
hy5cry1
hy5cry2
hy5cry1cry2

**Fig. S6 CRY1 seems to be main blue light photoreceptor upstream to HY5 in regulating iron uptake and root growth.** (a) Phenotype of WT, *hy5, cry1, cry2, cry1cry2, hy5cry1,hy5cry* and *hy5cry1cry2* plants grown on +Fe media for 5 days and then transferred to +Fe and -Fe media under blue light for 5 days. The double and triple mutants were screened on basis of hypocotyl length. Scale bar: 1 cm. (b) Boxplot of root length of WT, *hy5, cry1, cry2, cry1cry2, hy5cry1,hy5cry* and *hy5cry1cry2* plants grown on +Fe media for 5 days and then transferred to +Fe and -Fe media under blue light for 5 days. Means within each condition with the same letter are not significantly different according to one-way ANOVA followed by post hoc Tukey test, $P < 0.05$. using GraphPad PRISM 10. (c) Perls' and Perls' DAB stained maturation zone of WT, *hy5, cry1, cry2, cry1cry2, hy5cry1,hy5cry* and *hy5cry1cry2* plants grown on +Fe media under blue light for 5 days. The double and triple mutants were screened on basis of hypocotyl length. Scale bar: 100 µM. (d) Shoot phenotype of WT, *hy5, cry1, cry2, cry1cry2, hy5cry1,hy5cry* and *hy5cry1cry2* plants grown on +Fe and -Fe media under blue for 10 days. Scale bar: 0.5 cm. (e) Percentage reduction in chlorophyll in shoots of WT, *hy5, cry1, cry2, cry1cry2, hy5cry1,hy5cry* and *hy5cry1cry2* plants grown on +Fe and -Fe media under blue light for 10 days. Each biological replicate consisted of 5 shoots. Error bars represent ±SEM. Different letters (a, b, c, d) indicate significant differences, determined by one-way ANOVA followed by a post hoc Tukey Test ($P \leq 0.05$) using GraphPad PRISM 10.

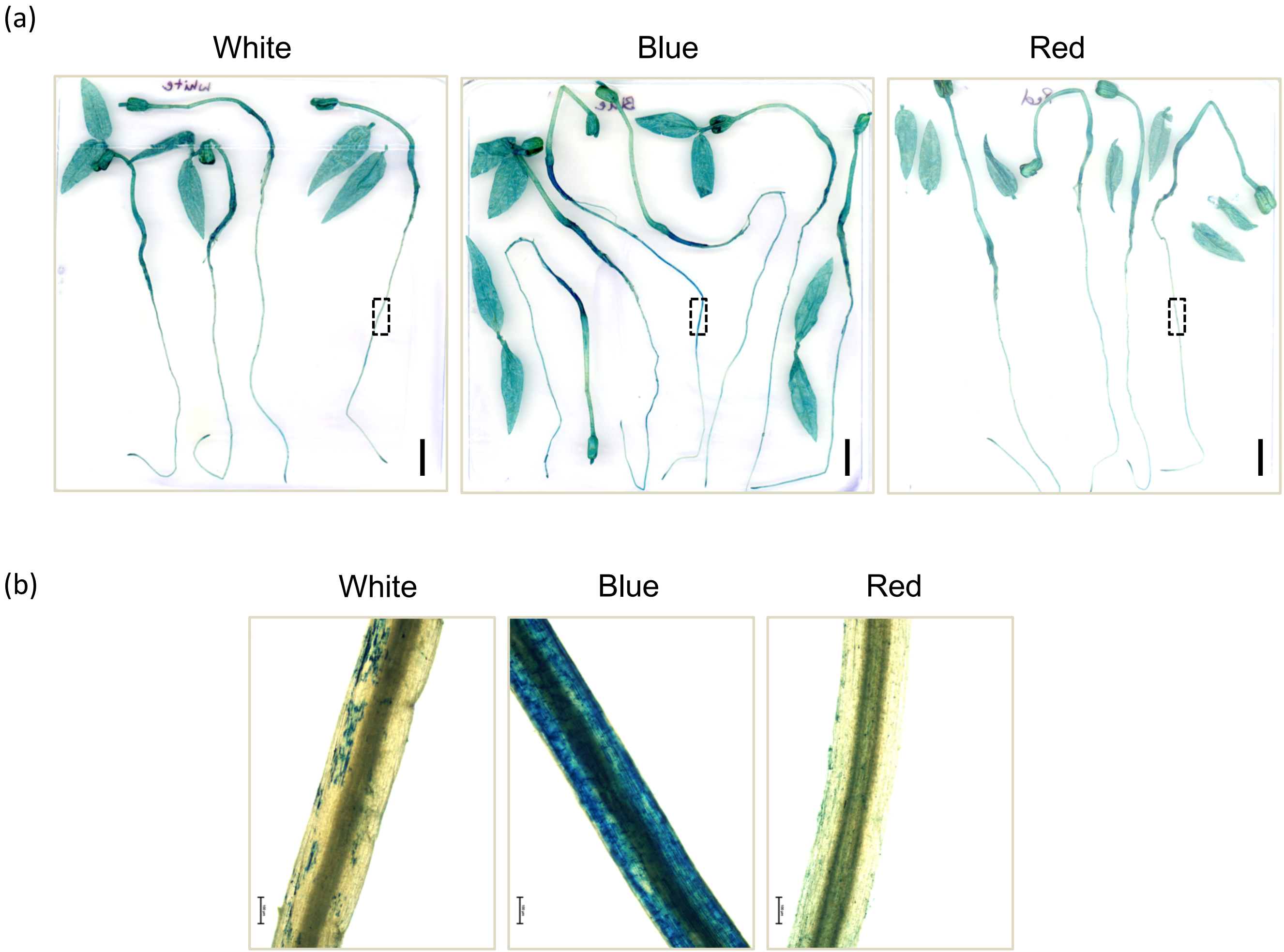


**Fig. S7 Moong bean sprouted in dark for 2 days were transferred to liquid Hoagland media for 5 days under different lights.** (a) Perls' stained seedlings of Mung bean , sprouted in dark for 2 days and then transferred to Fe-sufficient liquid Hoagland media under different lights for 5 days. Scale bar: 1 cm. (b) Perls' stained maturation zone of Mung bean marked in (a) as viewed under microscope. Scale bar: 100 µM.

**Table 1 Primers used in this study.**

| | |
|---|---|
| *hy5* LP | TTCACTCTCGATATCCGTTCG |
| *hy5* RP | ATGCGAGTGAATGACCATTTC |
| q*IRT1* FP | GAATGTGGAAGCGAGTCAGCGA |
| q*IRT1* RP | GATCCCGGAGGCGAAACACTTA |
| q*FRO2* FP | GCCACATCTGCGTATCAAGTT |
| q*FRO2* RP | TCCCAAACAAGCTACGACCA |
| q*HY5* FP | GAGGAGAAGCTGTCGGAAAA |
| qHY5 RP | CTCTGTTTTCCAACTCGCTCA |
| q*TUB* RP | CGACAATGAAGCTCTCTACGA |
| q*TUB* RP | AAGTCACACCGCTCATTGTT |
